# SysQuan: repurposing SILAC mice for the affordable absolute quantitation of the human proteome

**DOI:** 10.1101/2024.11.05.622109

**Authors:** Yassene Mohammed, Vincent R. Richard, M. Immanuel Reyes Madlangsakay, Ying Lao, Victor Spicer, Robert Popp, Claudia Gaither, Laura Henneken, Wolfgang Kleinekofort, René P. Zahedi, Christoph H. Borchers

## Abstract

Relative quantitation, used by most MS-based proteomics laboratories to determine protein *fold-changes*, requires samples being processed and analyzed together for best comparability through minimizing batch differences. This limits the adoption of MS-based proteomics in population-wide studies, and the detection of subtle but relevant changes in heterogeneous samples. Absolute quantitation circumvents these limitations and enables comparison of results across laboratories, studies, and longitudinally. However, high costs of the essential stable isotope labeled (SIL) standards prevents widespread access and limits the number of quantifiable proteins.

Our new approach, called “SysQuan”, repurposes SILAC mouse tissues/biofluids as system-wide internal standards for matched human samples to enable absolute quantitation of, theoretically, two-thirds of the human proteome using 157,086 shared tryptic peptides. We demonstrate that SysQuan enables quantification of 70% and 31% of the liver and plasma proteomes, respectively. We demonstrate for 14 metabolic proteins that abundant SIL mouse tissues enable cost-effective reverse absolute quantitation in, theoretically, 1000s of human samples. Moreover, 10,000s of light/heavy doublets in untargeted SysQuan datasets enable unique post-acquisition absolute quantitation.

SysQuan empowers researchers to replace relative quantitation with affordable absolute quantitation at scale, making data comparable across laboratories, diseases and tissues, enabling completely novel study designs and increasing reusability of data in repositories.

**TOC Graphic:** 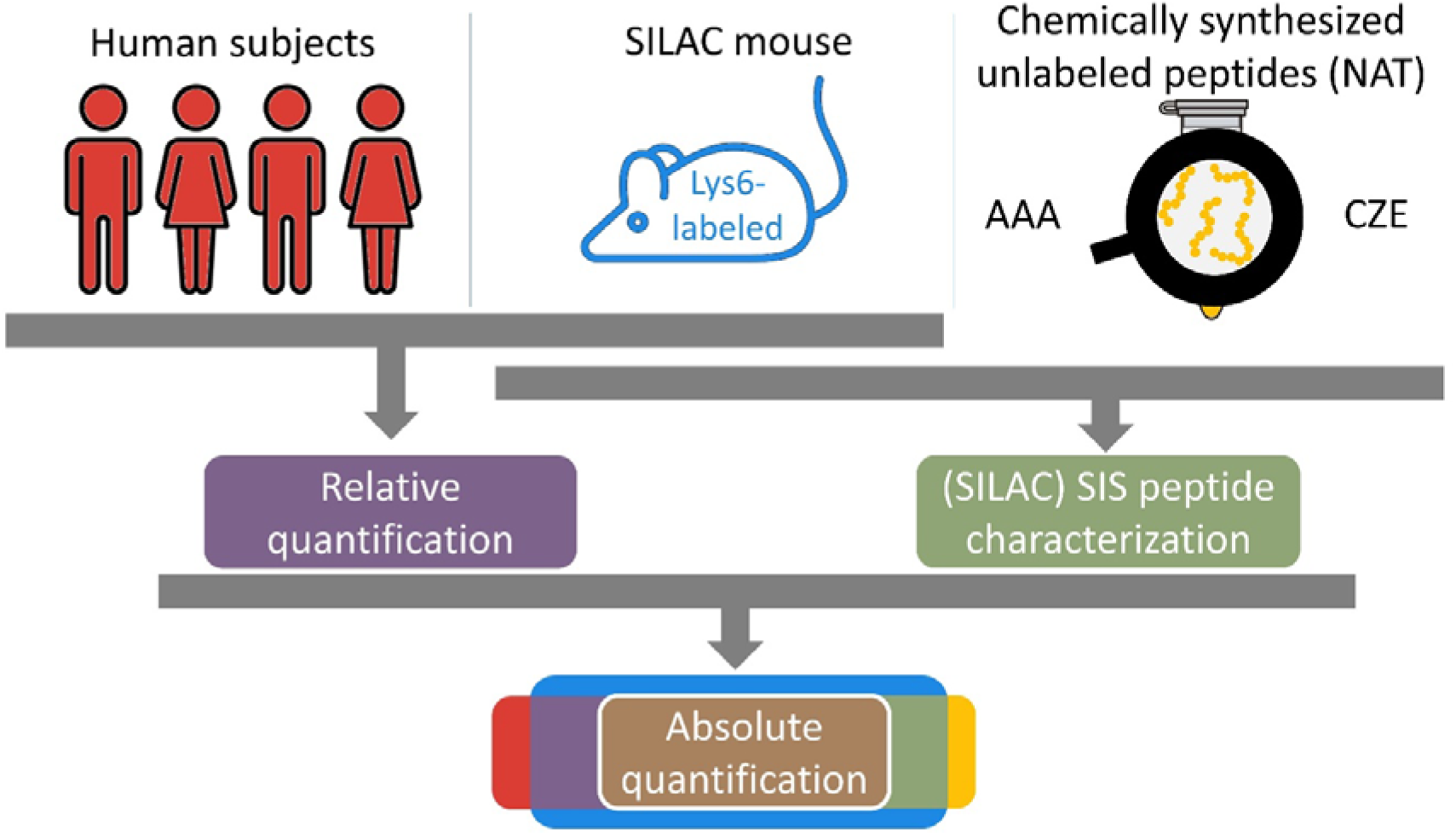

## Introduction

Proteins play a crucial role in health and disease and comprise the majority of disease biomarkers and drug targets. Mass spectrometry (MS) enables proteome-wide identification and (relative) quantitation of proteins with high sensitivity, precision, and unmatched specificity in virtually any type of sample, from mummies to human tissue biopsies. Consequently, MS-based proteomics has become the most important tool for biological discovery, with applications ranging from systems biology to biomarker candidate discovery.

Although, in recent years throughput and depth of MS-based proteomics approaches have considerably increased while concurrently cost per sample has been reduced (1), (pre)clinical studies continue to rely mainly on genomics approaches to study larger patient sample cohorts. One major reason why the impact of MS-based proteomics on large-scale studies is clearly lagging behind its unmatched potential to reflect actual disease phenotypes is that most laboratories exclusively employ *relative* quantitation to determine *fold-changes* in protein abundance between samples. This is because of ease of use of relative quantification methods, but also due to it being cost-effective. However, relative quantitation works best for a limited number of samples, which ideally are processed and analyzed together on the same LC-MS system. Once the number of samples increases and sample collection, preparation, measurement batches are introduced relative quantification suffers from considerable batch-effects that contradict the goals of low-abundance biomarker detection and target identification in heterogeneous patient populations.

Large-scale studies with 100s to 1000s of samples require quantitative reproducibility, that can eventually cover data generation across different laboratories and instruments as well as longitudinal measurements. To date, these requirements can *only* be met by absolute quantitation where protein concentrations are determined in individual samples using stable isotope labeled (SIL) internal standards (2). Spiking defined amounts of SIL peptides into samples enables quality control of the entire analytical workflow, including monitoring changes in instrument performance, and thus determining protein concentrations with maximum specificity and precision that can be compared across laboratories and platforms, between studies, and over time (3).

Targeted MS using multiple reaction monitoring (MRM) – also known as selected reaction monitoring (SRM) – on triple-quadrupole mass spectrometers is the “gold standard” for absolute quantitation, particularly in clinical settings (4, 5). However, having at least one SIL peptide for each target protein that is individually synthesized, purified, and characterized (for purity and quantity), drastically increases cost. The high cost of SIL peptides confine this powerful technology to a few applications and a few expert labs that mostly measure small protein panels, even though the feasibility of precise quantitation of 100s of proteins using highly multiplexed MRM has been demonstrated (6, 7). Consequently, MS-based proteomics is clearly dominated by more cost-efficient relative quantitation, which, apart from the abovementioned limitations, also lacks the precision to identify small but biologically relevant changes in protein abundance.

We present SysQuan – a novel strategy for cost-effective system-wide absolute quantitation of human proteins that we anticipate to reshape quantitative proteomics field. SysQuan enables determining precise concentrations for 1000s of proteins by exploiting the genetic proximity between humans and mice, with 99% of all human genes having homologs in the mouse genome. Homologous proteins in mice and humans may not have identical amino acid sequences, but they often share identical sequence stretches that are essential to protein function. We hypothesized that this *proteomic proximity* enables replacing costly SIL peptides with system-wide internal standards derived from SIL tissues that can be extracted in large quantities from SILAC (stable isotope labeling by amino acids in cell culture) mice (8). Matched “light” human and “heavy” SIL mouse tissues can be pooled, and after co-digestion peptides that are shared between humans and mice will generate light/heavy peptide pairs with defined mass shifts (doublets), just like with spiked-in SIL peptides or in conventional SILAC experiments.

We believe that the approach we are presenting here with SysQuan as a proteome-wide internal standard paves the way to MS-based large-scale proteomics studies with their unmatched precision and specificity to complement genomics and address the genotype phenotype disconnect (9–11).

## Experimental Procedures

### Human and SILAC mouse materials

Human plasma was purchased from BioIVT (Westbury, NY, USA) and obtained in K_2_-EDTA vials and stored at −80 °C until use. Plasma samples were from healthy donors (ages 18 to 50), who provided informed consent. Fresh-frozen normal-adjacent liver tissue from an adult human donor with hepatocellular carcinoma (male, 53 years old) was also obtained from BioIVT. Informed consent for the use of the samples in research was provided by the patients. Isotopically labelled mouse plasma and fresh-frozen liver samples were acquired from Silantes GmbH (Munich, Germany), product numbers 252923900 and 252923905. The labeling was performed in C57BL6 mice fed ^13^C-lysine (K+6) containing chow over 6 generations and was confirmed by the vendor to have 97% incorporation of K+6 (Silantes GmbH).

### Plasma sample preparation

Human and mouse plasma samples were depleted of the top 14 most abundant plasma proteins using High Select Top14 abundant protein depletion columns (Thermo Fisher Scientific), in order to reduce the dynamic range of the plasma proteome, and maximize the depth of proteome coverage. Depleted mouse and human plasma samples were pooled 1:1 and subsequently diluted in protein denaturation buffer (5% SDS, 100 mM TRIS pH 8.5, 10 mM TCEP) and heated to 60°C for 30 minutes. Proteins were reduced in 10 mM tris(2-carboxyethyl)phosphine) (TCEP), alkylated with 20 mM iodoacetamide (IAA), and digested with trypsin (Sigma) at a 1:10 enzyme:substrate ratio (approximately 5 µg of trypsin to 50 µg of depleted plasma protein) at 37°C for 16 hours using S-TRAP micro cartridges (Protifi LLC). Proteolysis was stopped with formic acid (FA; 0.5% v/v) and the resulting tryptic peptides were lyophilized to dryness prior to offline fractionation (detailed below).

### Liver tissue homogenization

Human (JGH) and mouse (Silantes) fresh-frozen liver samples were cryohomogenized using a prechilled (liquid nitrogen) mortar-and-hammer pestle. Homogenates were transferred to protein low-binding Eppendorf tubes containing protein extraction buffer (5% SDS, 100 mM TRIS pH 8.5, 10 mM TCEP) and subjected to probe-based ultrasonication (Thermo Sonic Dismembrator) and heating to 95°C for 10 minutes. Liver lysates were clarified by centrifugation (21000 x g, 5 minutes) and approximately 5% of the sample was reserved for protein concentration determination by bicinchonic acid assay (Thermo/Pierce).

### Liver SysQuan / UofM

Light human and heavy mouse liver lysate were mixed 1:1 to generate a 100-µg human/mouse protein mix. Disulfide bonds were reduced in 10 mM DTT for 30 minutes at 60°C followed by alkylation in 30 mM IAA for 45 minutes at room temperature in the dark. Proteolytic digestion was performed using single-pot solid-phase-enhanced sample preparation (SP3) (12). Prior to SP3, two types of carboxylate-modified SeraMag Speed beads (GE Life Sciences) were combined 1:1 (v/v), rinsed, and reconstituted in water at a concentration of 20 μg solids/μL. Ten microliters of the prepared bead mix were added to the lysate and samples were adjusted to pH 7.0 using HEPES buffer. To promote proteins binding to the beads, ACN was added to a final concentration of 70% (v/v) and samples were incubated at room temperature on a tube rotator for 18 minutes. Subsequently, beads were immobilized on a magnetic rack for 1 minute. The supernatant was discarded, and the pellet was rinsed twice with 200 μL of 70% ethanol and once with 200 μL of 100% ACN while on the magnetic rack. The rinsed beads were resuspended in 115 μL of 50 mM HEPES buffer (pH 8.0) supplemented with trypsin/LysC (Promega) at an enzyme-to-protein ratio of 1:25 (w/w) and incubated for 16 hours at 37°C. Peptide concentration was determined using Pierce Quantitative Fluorometric Peptide Assay (Thermo Fisher Scientific).

#### Offline high pH reversed-phase fractionation

An Agilent 1100 series LC system with UV detector (214 nm) and a 1 mm × 100 mm Gemini C18, 5 μm column (Phenomenex, Torrance, CA) were used for reversed-phase fractionation at pH 10. Both eluents (a) water and b (1:9 water: ACN) contained 20 mM ammonium formate pH 10. A linear gradient from 0.1-34% eluent B in 100 min was delivered at flow rate of 150 μL/min. A 65-μg aliquot of the liver digest was injected and eighty 1-minute fractions were collected (between 10 and 90 minutes) and concatenated into 40 to provide optimal separation orthogonality. These fractions were lyophilized and resuspended in 0.1% FA.

#### LC-MS/MS

Per fraction, 1 μg of total peptide was analyzed on an Orbitrap Exploris 480 (Thermo Fisher Scientific, Bremen, Germany) coupled to an Easy-nLC 1000 (Thermo Fisher Scientific) equipped with a C18 (Luna C18(2), 3 μm particle size (Phenomenex, Torrance, CA)) column packed in-house in Pico-Frit (100 μm X 30 cm) capillaries (New Objective, Woburn, MA). Peptides were separated using a binary gradient with (a) 0.1% FA and (b) 0.1% (v/v) FA in 80% ACN (LC-MS grade), ramping from 0 -5% B over 3 minutes, 5 – 7 % B over 2 minutes, 7-25% B over 84 minutes, 25 – 60 % B over 15 minutes, 60 – 95% B over 1 minute at a flow rate of 300 nL/min. The Orbitrap Exploris 480 instrument was operated in data-dependent acquisition mode. Spray voltage was set to 2.4 kV, funnel RF level at 40, and the heated capillary at 275°C. Survey scans covering the m/z 380–1500 were acquired at a resolution of 90,000 (at m/z 200), with a maximum ion injection time of 50 ms, and a normalized automatic gain control (AGC) target of 300%. This was followed by MS2 acquisition at a resolution of 30,000, selected ions were fragmented at 30% normalized collision energy, with intensity threshold kept at 2e4. AGC target value for fragment spectra was set to 100%, with a maximum ion injection time set to auto and an isolation width set at 1.6 m/z. Dynamic exclusion of previously selected ions was enabled for 30 seconds, charge state filtering was limited to 2–6, peptide match was set to preferred, and isotope exclusion was on.

### Liver SysQuan / McGill

Fresh frozen human and mouse liver samples were cryo-homogenized using a pre-chilled mortar and hammer pestle. Homogenates were transferred to protein low binding Eppendorf tubes containing protein extraction buffer (5% SDS, 100 mM TRIS pH 8.5, 10 mM TCEP) and subjected to probe based ultrasonication (Thermo Sonic Dismembrator) and heated to 95°C for 10 minutes. Liver lysates were then clarified by centrifugation (21K x g, 5 minutes) and approximately 5% of the sample was reserved for protein concentration determination by bicinchonic acid assay. Protein extracts were reduced and alkylated with 10 mM TCEP and 40 mM IAA respectively and an equivalent of 200 micrograms of liver lysate was digested with trypsin (Sigma) at a 1:10 enzyme to substrate ratio at 37°C for 16 hours using S-TRAP micro cartridges (Protifi LLC). Proteolysis was stopped with formic acid (0.5% v/v) and resulting tryptic peptides were lyophilized to dryness prior to offline fractionation. 50 µg of resultant Protein digests were then pooled 1:1 (100 µg total) and subjected to offline high pH reversed phase separation.

#### Offline high pH reversed-phase fractionation

To further increase the depth of coverage, we performed offline high-pH reversed-phase fractionation using an Agilent 1290 fraction collection system. Peptides were separated using a Waters XBridge peptide C18 column (2.1 x 150 mm, 2.1 µm particle) and peptides were separated using a binary gradient of (a) 10 mM ammonium formate (pH 10) and (b) acetonitrile with 10 mM ammonium formate (ACN) from 0% to 80% B over 48 minutes at flow rate of 400 μL/min with fractions collected every 30 seconds. Wells corresponding to fractions separated by 24 minutes or 48 fractions (i.e., well positions 1+49, 2+50, 3+51, etc. from left to right) were pooled and vacuum concentrated for a total of 48 fractions.

#### LC-MS/MS

Fractions from offline high-pH separation were reconstituted in 0.1% formic acid and analyzed by data-dependent acquisition (DDA) using an EvoSep One LC system (30SPD, EV1137 column at 45°C) coupled to a Bruker timsTOF HT mass spectrometer. Briefly, samples were acquired in DDA mode using 10 parallel accumulation serial fragmentation (PASEF) ramps scanning between 100 – 1700 m/z (method cycle time was approximately 1.1 seconds) (13). The capillary voltage was set to 1.6 kV. +2 and +3 charged peptides were selected based on their ion mobility values (0.6 – 1.6 1/k0). Precursors selected for ms/ms were dynamically excluded for 0.4 minutes.

### Data processing and analysis for untargeted acquisition

For each individual sample set (plasma, liver McGill, liver UofM) all raw data was searched using MSFragger embedded within the Fragpipe (v21.1) interface (https://fragpipe.nesvilab.org/). Data was searched against the canonical human proteome downloaded from UniProtKB (UP000005640, downloaded May 2024 containing 20,467 protein sequences). The FASTA file was supplemented with reverse decoy sequences and common contaminants using Fragpipe. Searches were performed using default settings with labeled Lys (K+6. 020129 Da) set as a variable modification and including precursor quantitation based on SILAC with the following label types: light (K+0) and heavy (K+6.020129). Peptide and protein false discovery rates were set to ≤ 1%. Further processing and figure generation was performed in R using standard analysis and plotting functions.

### Absolute quantitation of metabolomic pathway proteins

S-TRAP-digests of light human and SIL mouse liver were mixed 1:1 (w/w) based on BCA protein concentrations. A calibration curve was generated with 0.1, 1, 10, 100, 1000 fmol, each, of the NAT peptides for the 14 target proteins ACO1, ASS1, COX7C, DLAT, GAPDH, GOT1, HADH, NDUFA4, NDUFS4, NDUFV1, NDUFV2, SLC25A3, SUCLA2, SUCLG1 (see also Table 1 and Supplemental Table S1) in constant SIL mouse liver background. Next, human/mouse liver mix and the calibration curve were measured by LC-MRM on an Agilent 6495C mass spectrometer on-line coupled to an Agilent 1290 HPLC. Peptides were separated on an Agilent Zorbax Eclipse Plus RP-UHPLC column (2.1 x 150 mm, 1.8 µm) using a 47-min binary gradient (A: 0.1% FA; B: 100% ACN) ramping from 2-7% B in 2 min, 7%-27.6% B in 43 min, and 27.6-80% B in 2 min. A total of 930 MRM transitions were acquired for the 14 light and 14 SIL peptides using dynamic MRM and 2-min retention time windows, with minimum and maximum dwell times of 1.62 ms and 71.85 ms, respectively; the maximum cycle time was 555.59 ms (see Supplemental Data Table S2). Data was analyzed using Skyline. Calibration curves were used to calculate target analyte concentrations (fmol analyte / µg of liver tissue) in the human liver sample for a given light:heavy peak area ratio (see Supplemental Data Figure S1).

**Table 1:**
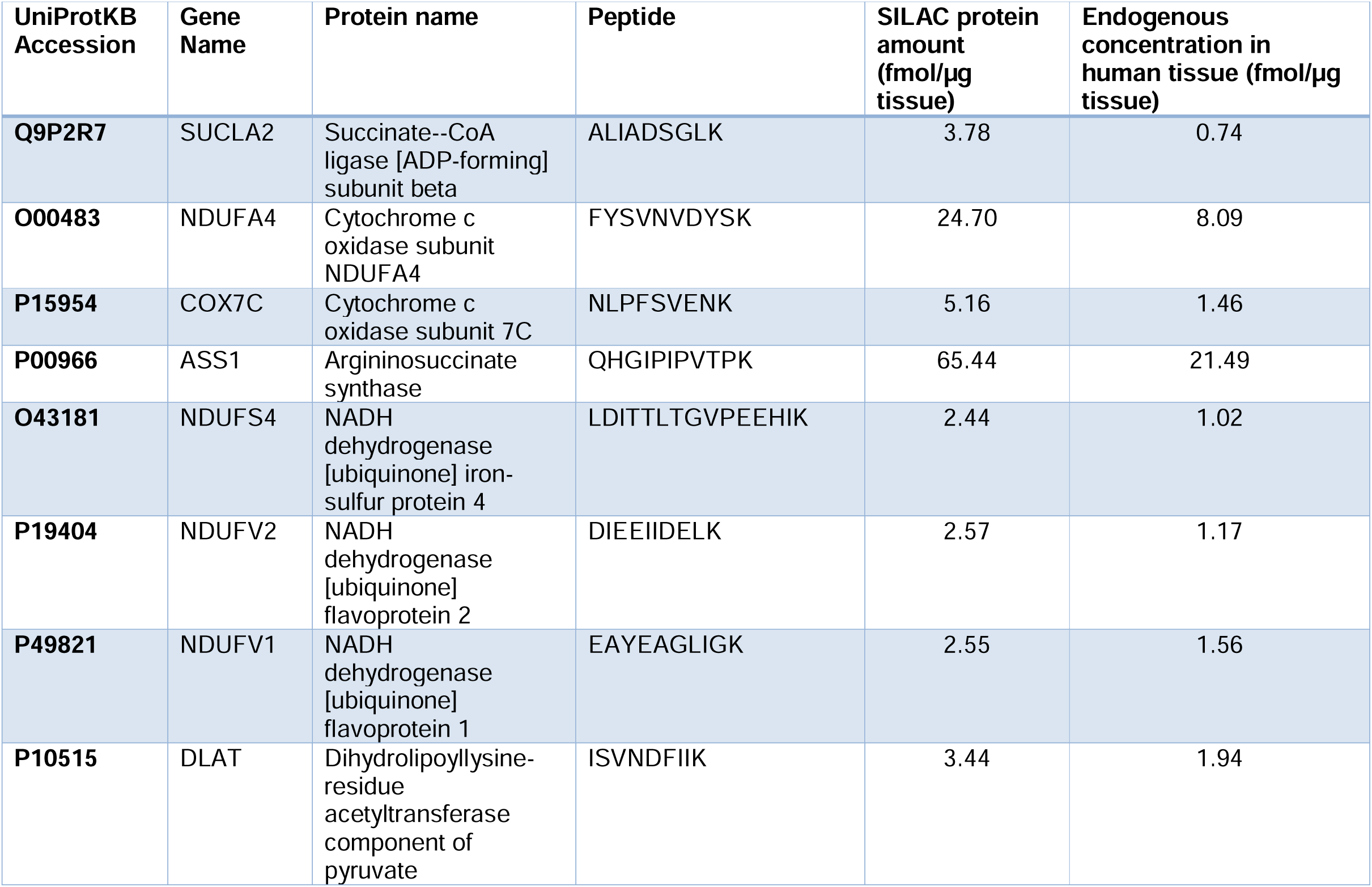
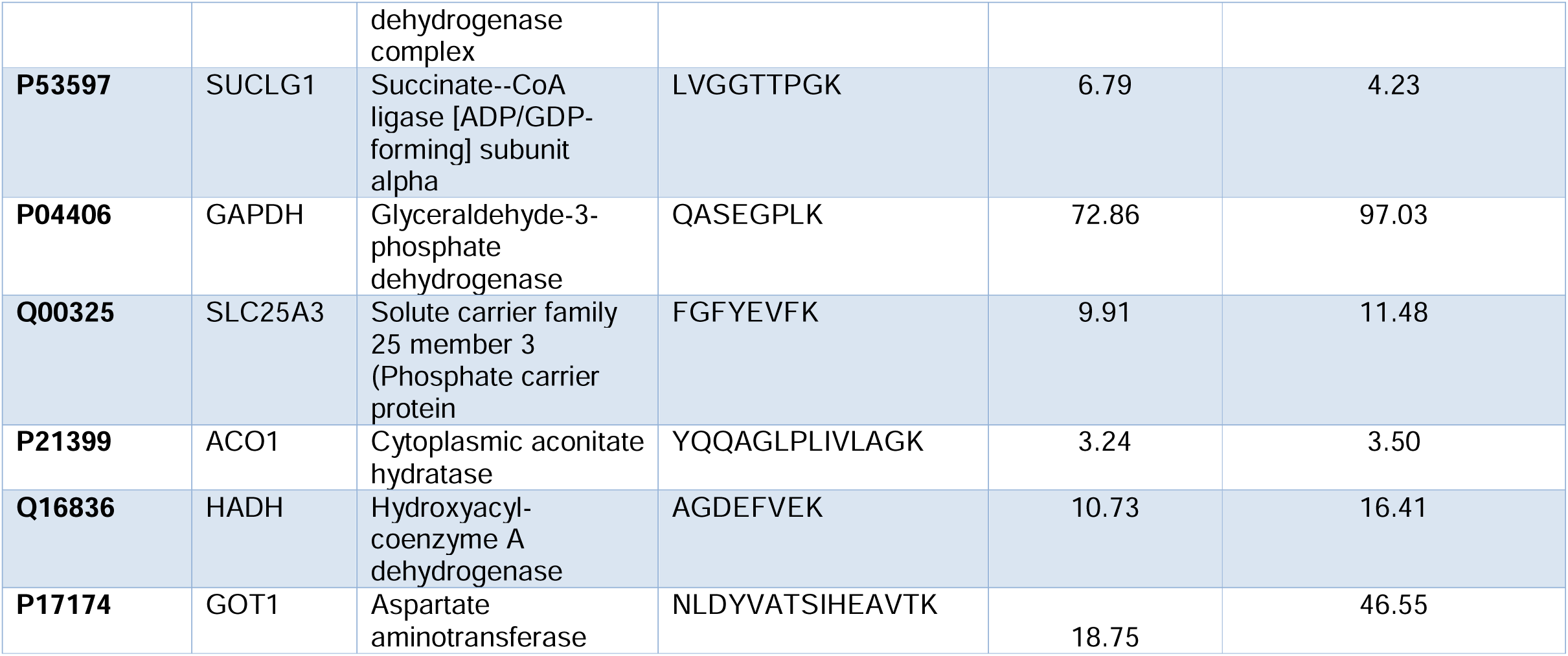
Absolute quantitation of metabolomic pathway protein components.

| UniProtKB Accession | Gene Name | Protein name | Peptide | SILAC protein amount (fmol/μg tissue) | Endogenous concentration in human tissue (fmol/μg tissue) |
| --- | --- | --- | --- | --- | --- |
| **Q9P2R7** | SUCLA2 | Succinate--CoA ligase [ADP-forming] subunit beta | ALIADSGLK | 3.78 | 0.74 |
| **O00483** | NDUFA4 | Cytochrome c oxidase subunit NDUFA4 | FYSVNVDYSK | 24.70 | 8.09 |
| **P15954** | COX7C | Cytochrome c oxidase subunit 7C | NLPFSVENK | 5.16 | 1.46 |
| **P00966** | ASS1 | Argininosuccinate synthase | QHGIPIPVTPK | 65.44 | 21.49 |
| **O43181** | NDUFS4 | NADH dehydrogenase [ubiquinone] iron-sulfur protein 4 | LDITTLTGVP EEHIK | 2.44 | 1.02 |
| **P19404** | NDUFV2 | NADH dehydrogenase [ubiquinone] flavoprotein 2 | DIEEIIDELK | 2.57 | 1.17 |
| **P49821** | NDUFV1 | NADH dehydrogenase [ubiquinone] flavoprotein 1 | EAYEAGLIGK | 2.55 | 1.56 |
| **P10515** | DLAT | Dihydrolipoyllysine-residue acetyltransferase component of pyruvate dehydrogenase complex | ISVNDFIIK | 3.44 | 1.94 |
| **P53597** | SUCLG1 | Succinate--CoA ligase [ADP/GDP-forming] subunit alpha | LVGGTTPGK | 6.79 | 4.23 |
| **P04406** | GAPDH | Glyceraldehyde-3-phosphate dehydrogenase | QASEGPLK | 72.86 | 97.03 |
| **Q00325** | SLC25A3 | Solute carrier family 25 member 3 (Phosphate carrier protein) | FGFYEVFK | 9.91 | 11.48 |
| **P21399** | ACO1 | Cytoplasmic aconitate hydratase | YQQAGLPLIVLAGK | 3.24 | 3.50 |
| **Q16836** | HADH | Hydroxyacyl-coenzyme A dehydrogenase | AGDEFVEK | 10.73 | 16.41 |
| **P17174** | GOT1 | Aspartate aminotransferase | NLDYVATSIHEAVTK | 18.75 | 46.55 |

## Results

Over the past 10 years, transformative advancements in MS-based proteomics have led to a significant boost in acquisition speed at improved proteome coverage, accompanied by a reduction in both required sample input and cost per sample. While in the past the considerable increase in sample complexity arising from mixing mouse and human proteomes had been a major challenge, studies involving ‘mixed proteomes’ such as PDX models (14, 15) and metaproteomics (16) studies with dozens of microorganisms have become feasible. It can be expected that the remarkable progress in proteomics technologies is going to continue, increasingly driven by single cell proteomics with its huge demands for higher throughput, greater sensitivity (17, 18), and lower cost per sample. Thus, to evaluate the future potential of SysQuan as proteome-wide internal standard, we performed in-depth proteomics profiling of pooled human/mouse liver tissue and plasma after extensive pre-fractionation (see Figure 1). As recent reports indicate that a novel hybrid mass spectrometer enables the analysis of 5000-6000 proteins in below 5 minutes (19), we believe that a similar coverage as achieved here with fractionation will be feasible 50-100x faster in single shot experiments.

**Figure 1:**
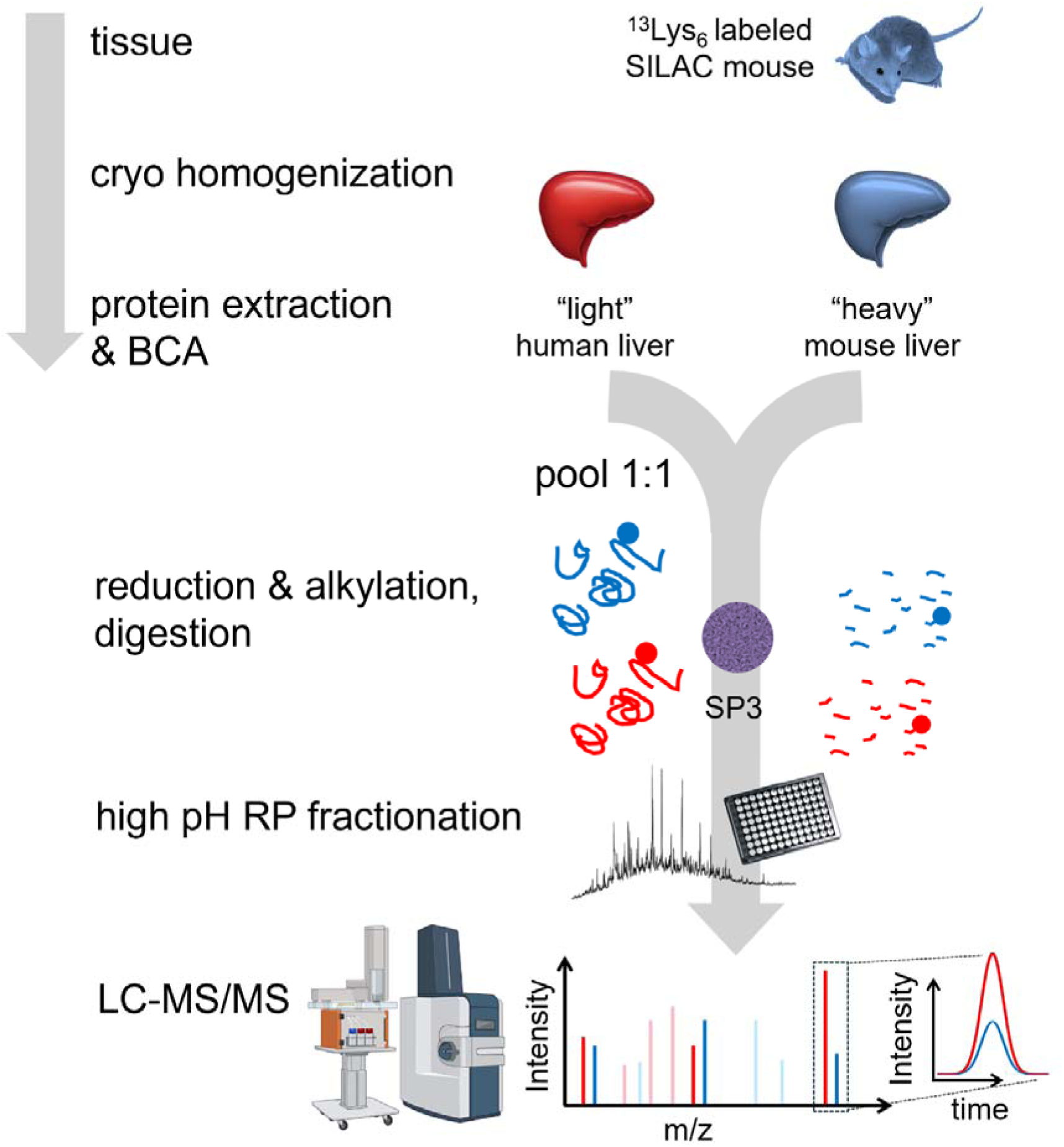
SysQuan feasibility workflow exemplified for liver tissue. Heavy mouse (blue) and light human (red) liver tissue were homogenized, proteins extracted, and pooled 1:1 (wt/wt). Proteolytic digestion with trypsin was performed using SP3, followed by offline reversed phase fractionation at pH 10.0. Fractions were analyzed by LC-MS/MS and light/heavy doublets representing SysQuan-quantifiable peptides were identified.

**Figure 2:**
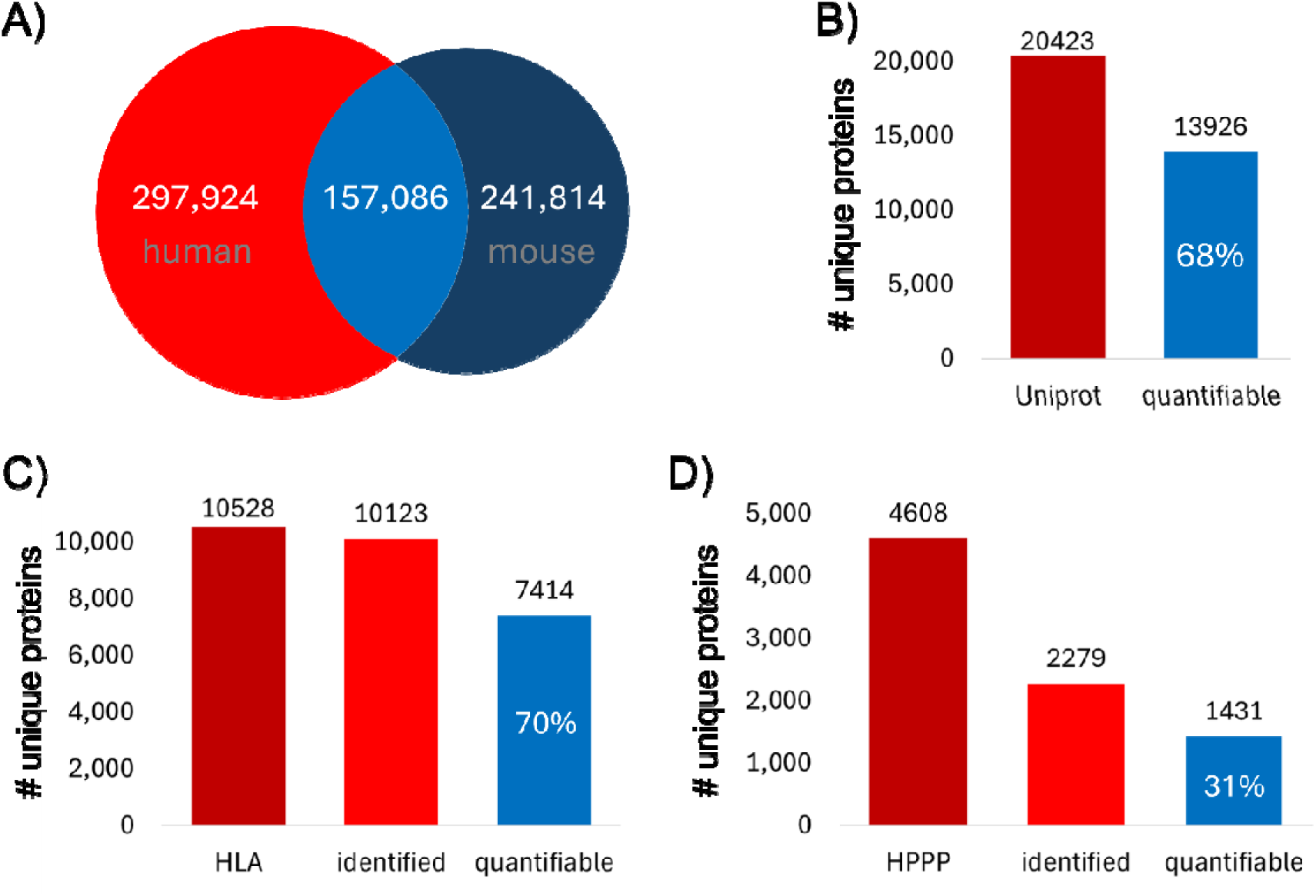
SysQuan human proteome coverage. (A) Overlap of *in-silico*-generated tryptic human and mouse peptides. (B) Based in this overlap, 68% of all human proteins in Uniprot are theoretically quantifiable using SysQuan. (C) Coverage and quantifiable share of the Human Liver Atlas (HLA), and (D) the Human Plasma Proteome Project (HPP) reference proteomes.

### Human-mouse proteomic homology

*Mus musculus* has high similarity with humans at the molecular level with 99% of human genes having homologs in the mouse genome. Further, many proteins in human and mouse share large part of their sequences. We calculated that around 68% of all human proteins have shared tryptic peptides in mouse proteome. Further, around 157k out of the 455k human tryptic peptides are also present in mouse, considering 7-25 amino acid length. SysQuan exploits this overlap by using SILAC mouse bio-fluids and tissues to quantify human.

### The SysQuan-quantifable proteome: liver

Mouse and human liver tissues were cryogenically homogenized, proteins extracted, and heavy mouse and light human protein extracts were mixed 1:1 (wt/wt), followed by reduction and alkylation, and proteolytic digestion with trypsin. The mouse/human peptide mix was then fractionated using high pH reversed phase chromatography. This workflow was performed independently with slight deviations at two different laboratories where individual fractions were analyzed by either DDA on an Orbitrap Exploris 480 (40 fractions) or PASEF-DDA on a timsTOF HT (48 fractions). Both datasets were processed with identical workflows using the Fragpipe software suite and SILAC settings.

In total we identified 59,734 and 115,762 peptide-spectrum matches (PSM) in the Exploris and timsTOF datasets, respectively, corresponding to 50,494 and 76,049 unique human peptide sequences, and 8323 and 8035 proteins. In the Exploris data, approximately 50% (23,727) of the unique peptides had light/heavy pairs corresponding to 5,990 human proteins, and 19,004 non-modified mouse/human peptide pairs corresponded to 5,560 unique proteins.

In the timsTOF data, 2/3 (48,760) of the unique peptides had light/heavy pairs, covering 6,987 unique human proteins, and 35,226 non-modified mouse/human peptide pairs corresponded to 6,305 unique proteins.

Focussing on one of the main liver function, metabolism, out of the 1562 proteins involved in the metabolic pathways (20), 1187 were identified in the liver proteome that we acquired, of which 1002 were quantifiable with heavy/light pairs.

### The SysQuan-quantifable proteome: plasma

Mouse and human plasma were depleted of the top 14 most abundant proteins, pooled 1:1, and processed as described for the liver/McGill dataset. A total of 48 high pH reversed phase fractions were analyzed by PASEF-DDA on a timsTOF HT. Fragpipe with SILAC settings yielded 23,501 PSM corresponding to 15,302 unique peptides and 2,279 unique proteins. More than 1/3 (5,769) peptides corresponding to 1,431 proteins had light/heavy pairs and 4,020 non-modified mouse/human peptide pairs corresponded to 1,209 proteins.

### Comparison to reference proteomes

To contextualize the depth of our SysQuan-quantifiable proteomes, we compared our data with the current reference proteomes from the proteomics community. The latest build of the Human Plasma Proteome Project (HPPP 2023-04) of the Human Proteome Organization,(15) combines results from 313 experiments and comprises 4,608 human plasma proteins. In our mouse/human plasma mix we identified 2,279 human proteins, 1431 of which can be quantified using SysQuan. Thus, with a single workflow and proteolytic enzyme, SysQuan enables quantitation of approximately 31% of the known plasma proteome (see Table 2). Diversifying the proteomics workflow to better reflect the 313 experiments in HPPP may allow quantifying up to 2/3 of the plasma proteome in accordance with our *in silico* predictions. Integrating SysQuan into recent strategies that have considerably increased the depth of plasma proteomics (21, 22) may help reaching this ambitious goal.

The human liver atlas contains 10,528 unique proteins,(23) identified through the use of diverse immortalized liver cell lines, primary cells, and human biopsies, amounting to a total of 34 different sample types. Using mixed human/mouse liver samples, we were able to identify 96% of the known human liver proteome, and our data indicate that 70% of this known liver proteome is quantifiable using SysQuan, which aligns well with our *in silico* predictions.

### Labeling efficiency

Krüger et al reported average SILAC mouse labeling efficiencies of >95% after four generations of labeling (8). To determine our labeling efficiency, we analyzed the SILAC mouse liver as described above for the McGill mouse/human mixes after extensive prefractionation (48 fractions, tims TOF HT). For each identified mouse peptide, we used the SILAC light and heavy channel intensities to determine the % labeling efficiency. The average labeling efficiency over 70,287 peptides representing 7,703 mouse proteins was 96.0%, which is well in-line the data from Krüger et al. Even for synthetic SIL peptides, a labeling efficiency >95% is widely considered sufficient for absolute quantitation. Importantly, the knowledge on the exact labeling efficiency for individual mouse peptides in a given SysQuan tissue can be leveraged to correct human protein concentrations and further increase quantitative accuracy – similar to correction factors for isobaric stable isotope labeling methods.

### Translating SysQuan to targeted proteomics

We envision that once mouse protein concentrations have been determined, SysQuan enables proteome-wide absolute quantitation of human proteins. While non-targeted proteomics methods enable in-depth proteome analyses, targeted MS with scheduling and SIL peptides has been used for absolute quantitation of 100s of proteins in a single LC-MS run with unmatched precision, dynamic range, and sensitivity. Targeted MS can provide access to proteins that are below the limit of detection/quantitation of non-targeted methods.

To demonstrate the potential of SysQuan for absolute quantitation of human samples, we focused on a few liver proteins that map to metabolic pathways. From 1002 quantifiable proteins associated with various metabolic pathways, we selected 14 that cover various aspect parts of metabolism including electron transfer cycle, mitochondrial function, tricarboxylic acid (TCA) cycle, glycolysis, amino acid metabolism, beta-oxidation, and mitochondrial transport (Supplemental Data Table S1). These proteins included ACO1, ASS1, COX7C, DLAT, GAPDH, GOT1, HADH, NDUFA4, NDUFS4, NDUFV1, NDUFV2, SLC25A3, SUCLA2, and SUCLG1, for which we purchased light synthetic standard peptides (NAT; from MRM Proteomics Inc). NAT purity was determined using capillary zone electrophoresis (CZE) and NAT quantity was determined using amino acid analysis (AAA). For all 14 targets, MRM conditions were optimized on an Agilent 6495D using the inexpensive NAT peptides. Next, we mixed heavy mouse/light human liver digests 1:1 (w/w) and measured the corresponding heavy/light pairs by MRM, followed by a calibration curve with 0.1, 1, 10, 100, 1000 fmol of each NAT in a constant SIL mouse liver background. Light/heavy peak ratios were determined to calculate human liver protein concentrations in fmol per µg liver tissue for all 14 target proteins (see Table 1), covering more than two orders of magnitude in dynamic range.

Next, we compared these results of the determined human liver protein absolute concentrations with data from proteomicsDB.org (24), where protein abundance in individual cells or tissues is *estimated* based on intensity-based absolute quantitation (iBAQ) (25) distributions over dozens of datasets. Although not accurate and not absolute in terms of referencing to standards, such estimates of protein abundance based on spectral counts or intensities have proven useful to get an overall idea of the composition of a given proteome (26, 27). Interestingly, and despite the completely different analytical workflows, protein concentrations determined by SysQuan correlated well with iBAQ estimates from proteomicsDB.org, with an R^2^ of 0.89, which is an excellent confirmation of both methodologies (Figure 3).

**Figure 3:**
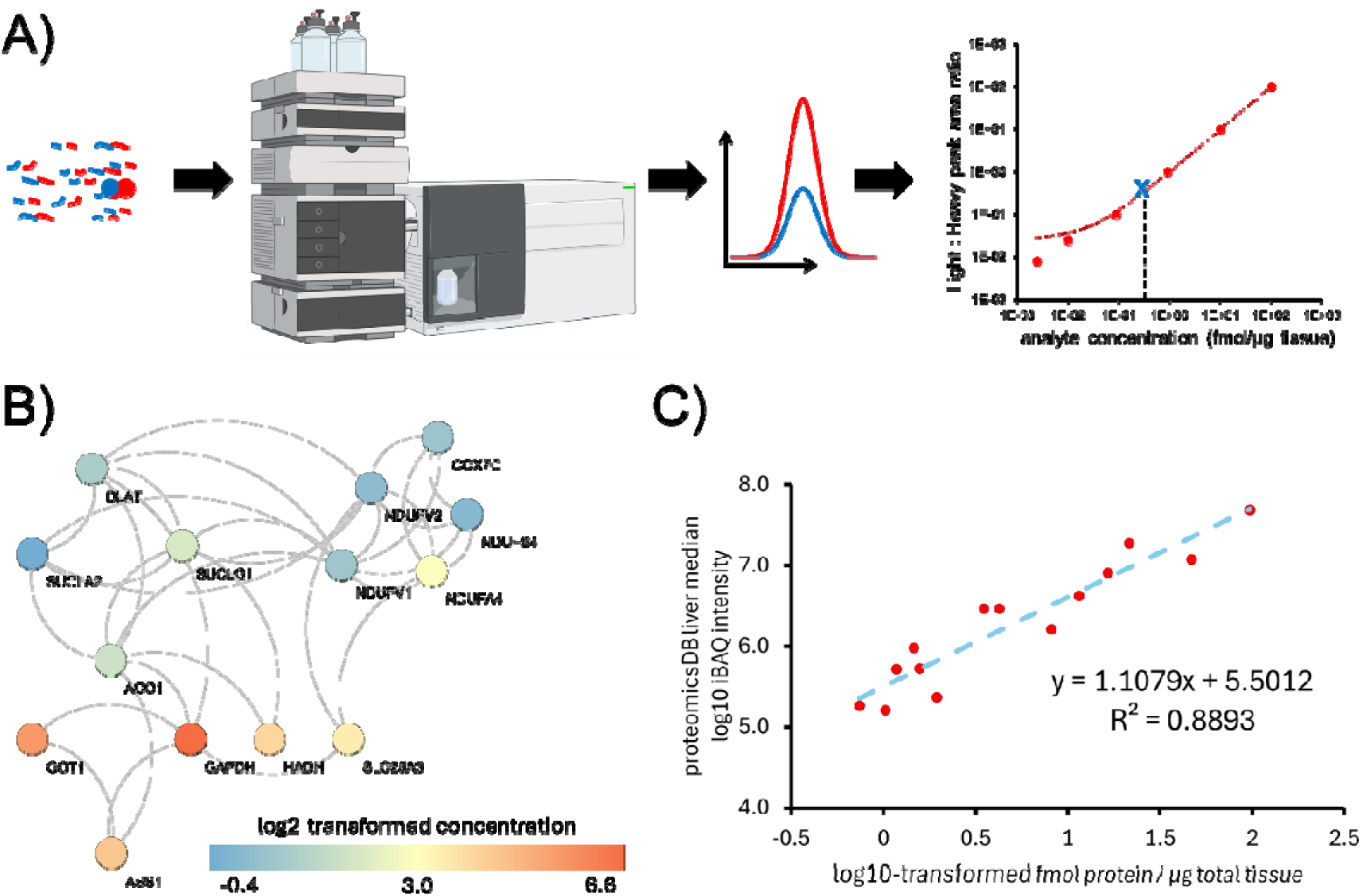
Absolute quantitation of metabolomic pathway proteins. (A) Overall workflow. (B) STRING protein/protein interaction network. (C) Correlation of SysQuan-derived concentrations and iBAQ-derived estimates for protein levels in proteomicsDB.

These results confirm that absolute quantitation using SysQuan is feasible. The same SILAC tissue reference standard can be used for characterizing 100s-1000s of samples, thus representing a cost-effective and concurrently more comprehensive alternative to current SIL peptide-based absolute quantitation. A recent study using SIL peptides and guided data independent acquisition parallel accumulation and serial fragmentation (g-dia-PASEF) reported absolute quantitation of 579 human proteins over a dynamic range of 4 orders of magnitude in depleted plasma samples (28). We have recently tested that state-of-the-art LC and triple quadrupole instrumentation allow reducing the run time for multiplexed MRM of plasma protein quantification by almost 4 times while maintaining depth and quantitative precision. Based on these results, we believe that the available LC-MS technology already enables quantifying 1000 proteins by targeted MS in 30-60 minutes (i.e., targeting both light and heavy forms of one peptide per protein and having a fairly even distribution of retention time). While this is not comparable to the proteome depth of non-targeted approaches, targeted MS provides improved precision (and accuracy) and unlocks access to a wider dynamic range, thus particularly enabling the quantitation of very low-abundance targets that are otherwise not accessible. This is particularly important as in many studies, precise measurement of a smaller but defined set of proteins across a large number of samples and without missing values is preferrable over proteome-wide analysis with less precision and more incomplete data.

Notably, SysQuan-based absolute quantitation can be transferred to non-targeted methods, such as DDA or DIA-based label free quantitation. Interestingly, using non-targeted SysQuan for absolute quantitation provides a unique feature: all light/heavy doublets present in an acquired dataset can be *retroactively* absolutely quantified once the concentrations of the same mouse reference standard have been determined, i.e., even years post-acquisition.

### Precision and accuracy

Quantifying proteins using proteotypic peptides as surrogates is widely accepted as precise, i.e. reproducible, and is the basis of relative quantitation in bottom-up proteomics. Achieving accuracy, i.e. determining protein concentrations that are close to the true value, however, is considerably more challenging and in the absence of reference values virtually not possible. It requires additional effort, including optimizing and determining digestion efficiency associated with the target proteotypic peptide release, as well as determining recovery of the protein and the surrogate peptides. While all of this can be easily performed in surrogate matrices, for example by spiking a recombinant protein in a matrix devoid of endogenous protein, it is difficult to perform in patient samples with often unknown levels of the protein-of-interest. Standard addition using certified authentic material of known concentration enables accurate quantitation,(29) but such certified materials are, unfortunately, not available for most proteins, and it is often impossible or impractical to obtain them (30).

What matters most in practice and for preclinical applications is precision, since it encompasses repeatability and reproducibility. For most applications, from discrimination between treatment groups to identification of biomarker candidates, precision is sufficient as it enables comparison of data, if internal standards are being used also across platforms and laboratories. Traditional targeted proteomics using internal standards provides the required precision, regardless of whether these internal standards are synthesized or SILAC mouse-derived SIL peptides. Notably, the use of SIL proteins to generate SIL peptide internal standards out of sequence stretches that are common between humans and mice, adds an inherent digestion control, addressing one of the main challenges of absolute quantitation using SIL peptides. This has been partially addressed by strategies such as QCoNCAT (31) where stable isotope labeled proteins are expressed and used as internal standards.

### Reproducibility in SysQuan

The use of SIL mouse tissues and biofluid as internal standard stocks improves precision of relative quantitation and enables comparing large sample numbers even measured in different labs, as human proteins can be referenced to the same internal standard. Once the concentrations of a set of mouse proteins are determined in a given reference SIL tissue stock, this standard enables absolute quantitation of those proteins across 10,000s of samples. Concurrently, this overcomes two major challenges that impair the precision and accuracy of current absolute quantitation methods: matrix effects and variability in digestion efficiency (10, 32), both of which cannot be controlled using SIL peptides (9).

To test this, we analyzed the liver samples at two different laboratories, with different sample preparation, different prefractionation procedures, different reverse LC systems and different mass spectrometers. We found a very good agreement of protein quantification between the two acquisitions with two main insights (Figure 4). First, in both measurement the log_2_ ratio between human and SILAC SIL mouse peptides were centered around zero, indicating that the concentration of the human proteins and released peptides by digestion are very similar to their SILAC SIL mouse counterparts. This reinforce the idea that matched SILAC mouse tissue can be used effectively to quantify human tissue “out-of-the-box” with minimal matrix effect. Alternative methods of generating SILAC SIL peptides in cell culture (33, 34) cannot account for this since the SILAC cells and actual samples have two completely different proteomes. Second, we assed the agreement between the two quantification measurements using Bland-Altman analysis (35). We calculated that 95.35% of all quantifications were within the upper and lower limit of agreement – defined by mean of quantification difference ± standard deviation of quantification difference. This robustness across different laboratory experimental conditions suggests that SysQuan quantification is reproducible, reliable, and independent of laboratory, sample preparation procedure, or instrumentation. This is a crucial aspect for translational research applications and should enable application of proteomics in longitudinal and population-wide studies.

**Figure 4.**
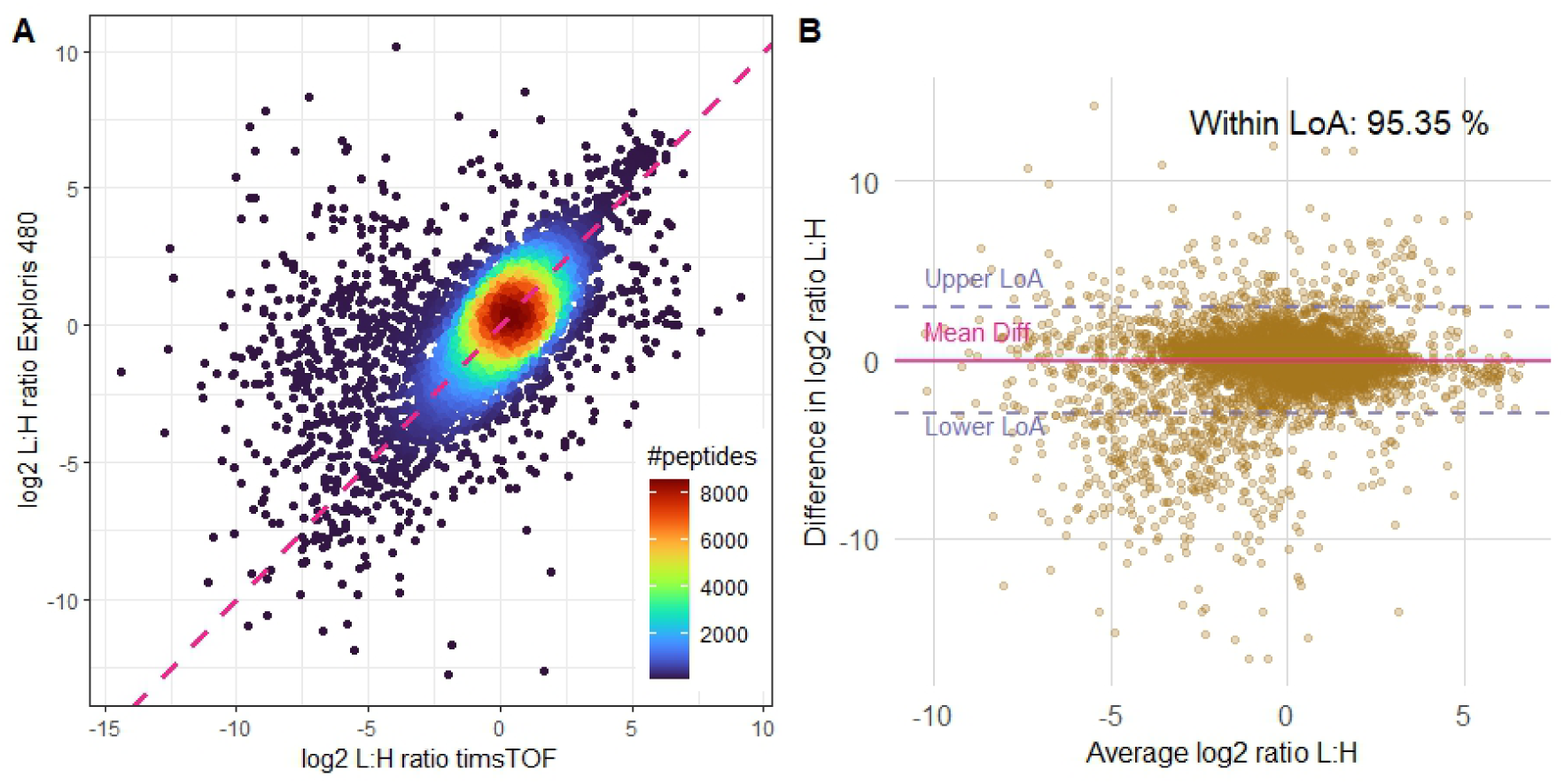
Agreement of quantification between two sites of measurement with different sample preparation and instrumentation. (A) regression between the two analysis on two different liquid chromatography systems as well as mass spectrometers showing that most of the Log_2_ ratios are centered around 0. (B) Bland-Altman analysis showing an agreement of 95.35% between two measurements. LoA: limit of agreement; Mean Diff: mean difference.

## Discussion

High-throughput molecular phenotyping methods, such as RNA sequencing, metabolomics, and proteomics, are the cornerstones of modern biological and clinical studies, providing crucial mechanistic insights at the molecular level. Despite technological advancements, however, a key challenge remains -- the affordability of these methods as precise and selective tools for determining and quantifying system-wide molecular changes. In contrast, the remarkable achievements in genomics can largely be attributed to efforts focused on making sequencing both rapid and reproducible at reduced cost.

Mass spectrometry-based proteomics has made significant strides in recent years, transitioning from qualitative to quantitative approaches. Label-free quantification techniques have emerged as cost-effective methods for biomarker discovery studies and hypothesis generation. However, despite the considerable increase in throughput accompanied by an increase in proteome coverage and concurrent decrease in cost per sample, experimental designs are increasingly facing constraints. These constraints arise from the need that samples need to be processed and analyzed together to avoid considerable batch effects that impair statistical analysis and consequently the identification of relevant biological changes. The only way to address these limitations is through determining the *amount* of a protein relative to a spiked-in internal standard, which allows subtle changes in protein levels can be detected. This enables more reliable comparisons between sample sets even across different batches and regardless of whether these batches have even been analyzed on the same platform. However, the high cost of chemical synthesis of heavy labelled internal standards becomes prohibitive when quantifying large numbers of proteins, creating a significant barrier to widespread adoption. SysQuan can overcome this barrier by making quantitative proteomics using internal standards – and even system-wide absolute quantitation – more accessible and affordable. This is exemplified by the high coverage of the plasma and liver proteomes achieved in this study. This coverage can be further increased by the use of alternative enzymes: in silico digestions with LysC and GluC indicate that an additional ∼1000 human proteins may be quantifiable compared to trypsin. Even the use of low-specificity enzymes such as subtilisin and thermolysin might be feasible (36), however, needs empirical testing as it is currently impossible to reliably predict their cleavage products.

Importantly, SysQuan also enables post-acquisition absolute quantitation from untargeted proteomics datasets. Using NAT peptides to *(i)* quantify SIL proteins in SILAC mouse tissue *(ii)* to be used as internal standard for reverse quantitation of human proteins in 100s of samples is significantly more cost-effective than the current gold-standard using synthetic SIL peptides (37).

We believe that SysQuan can lead to a paradigm shift, as it enables researchers to replace relative quantitation with affordable absolute quantitation at scale, thus making every single proteomics study and individual sample considerably more informative and valuable for the community. This will be particularly relevant in the study of rare diseases, typically suffering from limited availability of patient samples. The ability to directly compare individual patients and studies, even when single cases have been analyzed and published years earlier, will transform rare disease research. We envision a utility for SysQuan as companion diagnostic and even diagnostic tool, as beyond classical absolute quantitation of biomarkers it will provide direct and clear evidence for the dysregulation of key metabolic pathways – without the need to run patients samples in batches with controls. The ability to compare protein concentrations across different human tissues and different diseases will enable novel study designs that are impossible with current technology.

SysQuan can be used for both targeted proteomics (MRM and parallel reaction monitoring, PRM) and non-targeted proteomics (data dependent and data independent acquisition, DDA and DIA). SysQuan will, for the first time, enable researchers to retroactively quantify protein concentrations in DDA/DIA datasets, even if mouse SIL standard protein concentrations are determined years post-acquisition. This unique feature opens new avenues for the reanalysis and repurposing of existing datasets within the scientific community (11, 38).

Similarly, the capability to universally compare protein concentrations across datasets will drive entirely new, community-driven studies based on the reanalysis and repurposing of datasets in public repositories such as PRIDE (39) and MassIVE (40), further enhancing their value and impact on biomedical and fundamental research. The ability to retroactively determine protein concentrations in existing datasets, once the corresponding SIL mouse protein concentrations have been determined, opens up virtually unlimited possibilities for discovery.

In conclusion, we believe that SysQuan has the potential to transform fundamental and translational biomedical research, addressing the reproducibility crisis in life science (41) by making standardized quantitative proteomics broadly accessible (42).

## Acknowledgments

CHB is grateful for support from the Segal McGill Chair in Molecular Oncology at McGill University. CHB and RPZ are also grateful to Genome Canada and Genome Quebec for funding via the MutaQuant GAPP (#6567 Borchers_Zahedi_APF). CHB is grateful for support from the Warren Y. Soper Charitable Trust for the Warren Y. Soper Clinical Proteomics Centre at the Jewish General Hospital (Montréal, Quebec, Canada). CHB. is also grateful for support from the Terry Fox Research Institute and the Alvin Segal Family Foundation for the Segal Cancer Proteomics Centre at the Jewish General Hospital (Montréal, Quebec, Canada). RPZ is grateful for support from the University of Manitoba, Shared Health Manitoba, and support from the H. E. Seller Research Chair in Internal Medicine.

This work was done under the auspices of a Memorandum of Understanding between McGill and the U.S. National Cancer Institute’s International Cancer Proteogenome Consortium (ICPC). ICPC encourages international cooperation among institutions and nations in proteogenomic cancer research in which proteogenomic datasets are made available to the public. This work was also done in collaboration with the U.S. National Cancer Institute’s Clinical Proteomic Tumor Analysis Consortium (CPTAC).

Figure 1 and Figure 3 instrument cartoons were created in BioRender by Richard, V. (2024), https://BioRender.com/d21u030.

## Data Availability

The mass spectrometry proteomics data have been deposited to the ProteomeXchange Consortium via the PRIDE, MassIVE, and PanoramaWeb partner repositories. The plasma and liver data acquired at McGill are available in PRIDE under the dataset identifier PXD057266 (reviewer access details as follow; Project accession: PXD057266; Token: DJrJ58c7LVk3) as well as PanoramaWeb available here: https://panoramaweb.org/jJdH00.url. Alternatively, reviewer can access the dataset by logging in to the PRIDE website using the following account details; Username, Password: wEYzqDyRmzqB).

The liver data acquired at UofM is available in MassIVE under the dataset identifier MSV000096198 (reviewer access with password: cNfyQ5HGb4u1bIkR).

## Supplemental Data

**Supplemental Data Figure S1.**
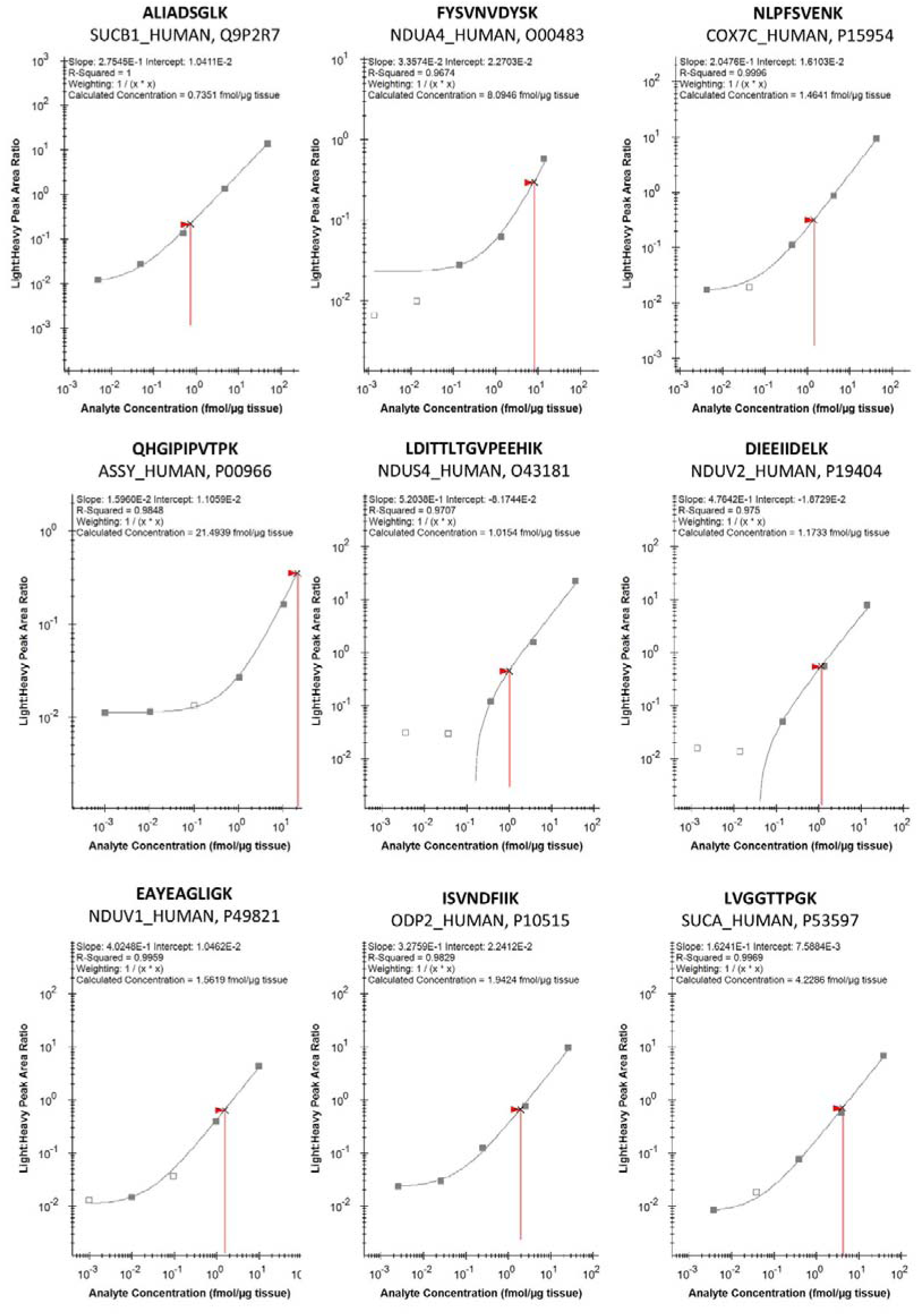
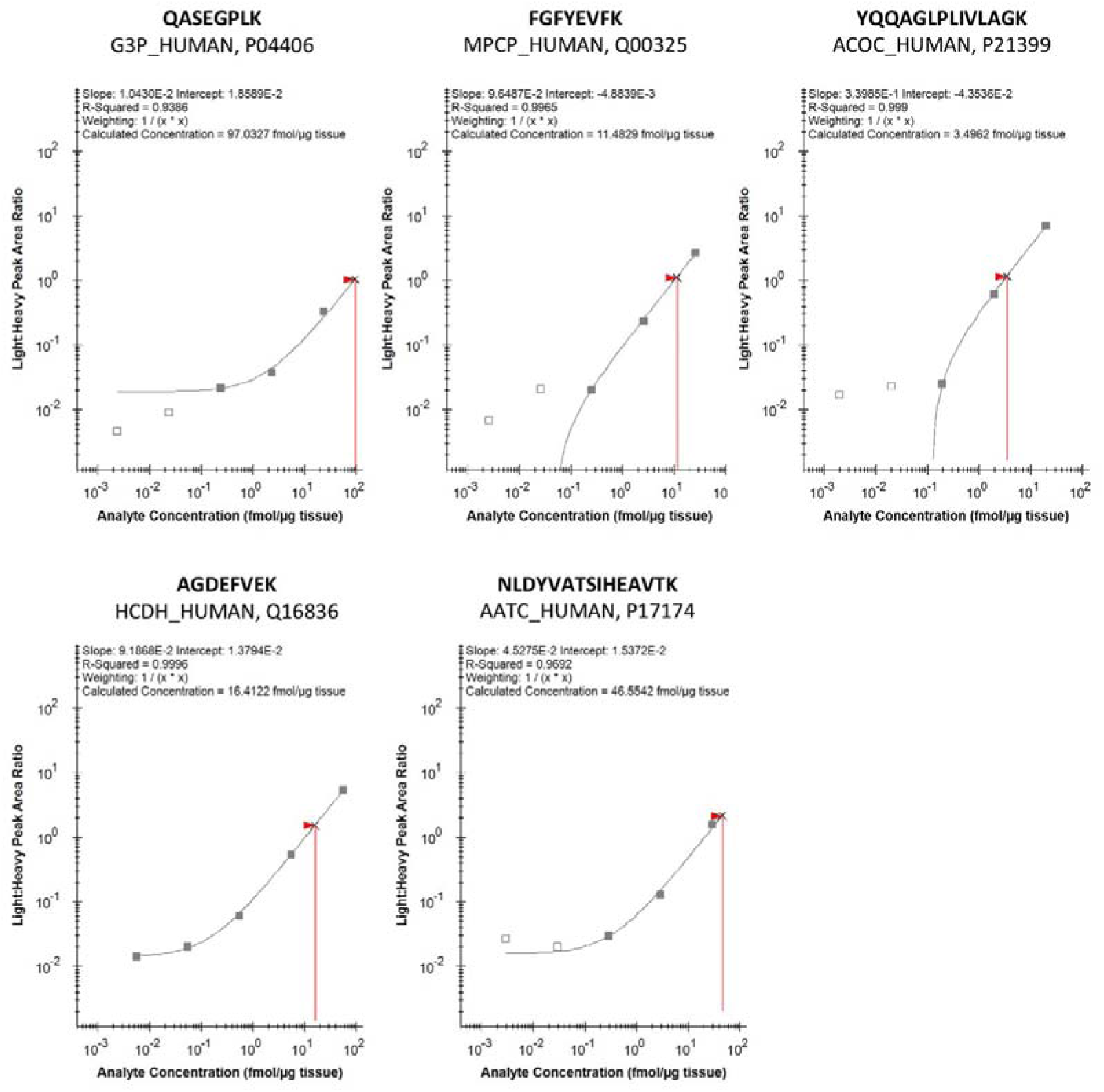
Calibration curves to quantify 14 proteins associated with metabolic pathways

**Supplemental Data Table S1.**
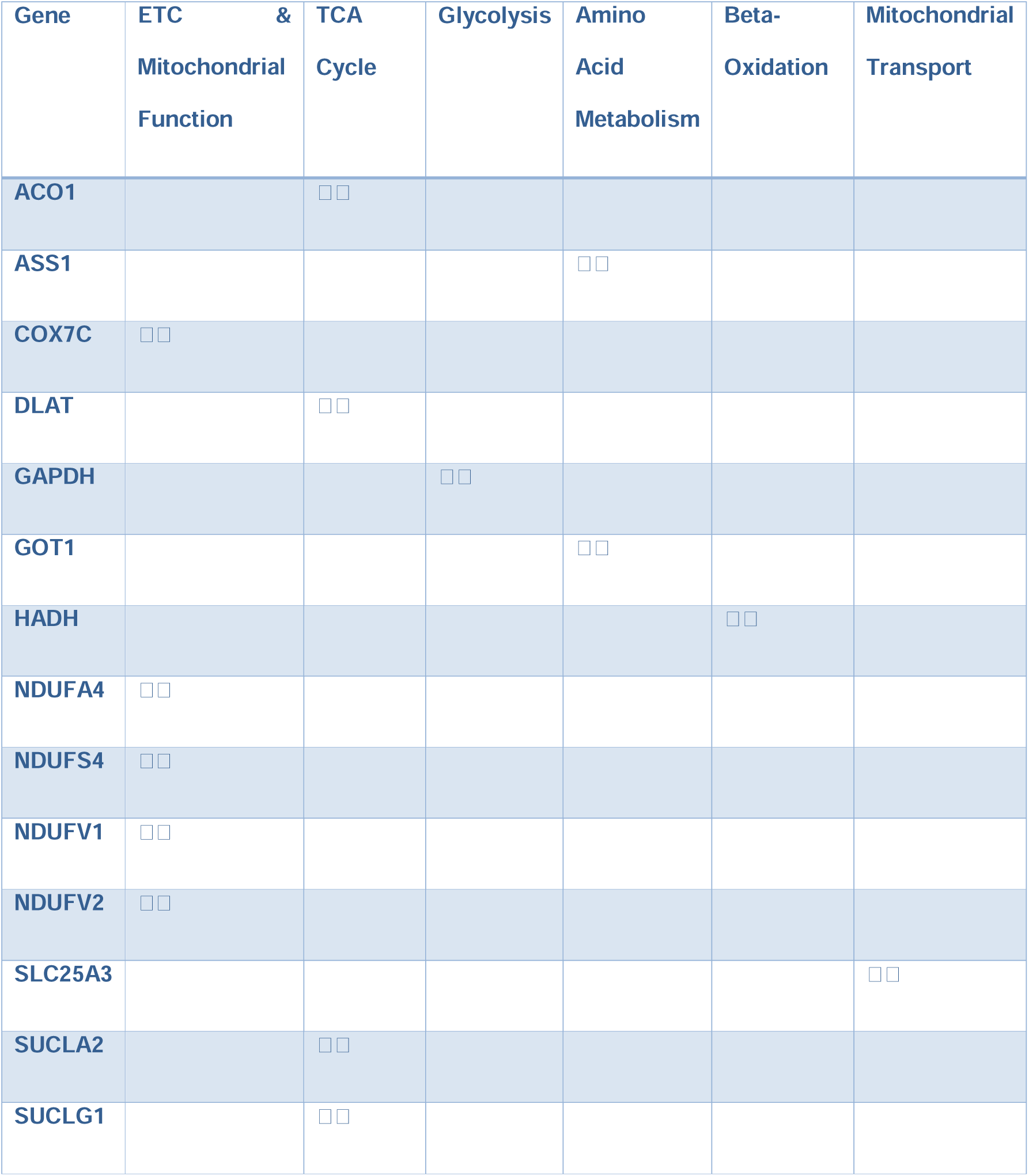
Proteins associated with various metabolic pathways for which we performed absolute quantification in human liver by double standards SILAC SIL and synthetic NAT.

**Supplemental Data Table S2.**
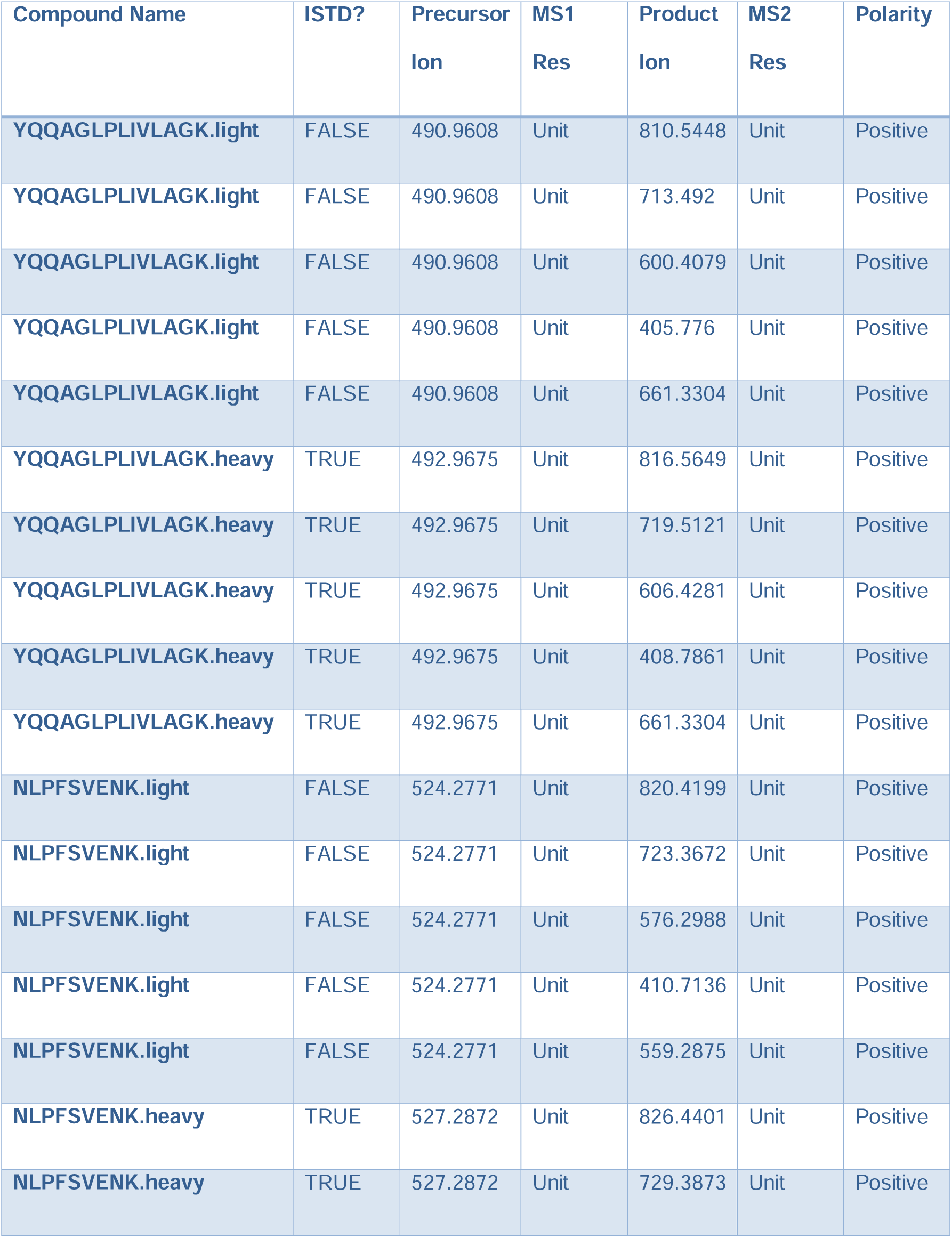
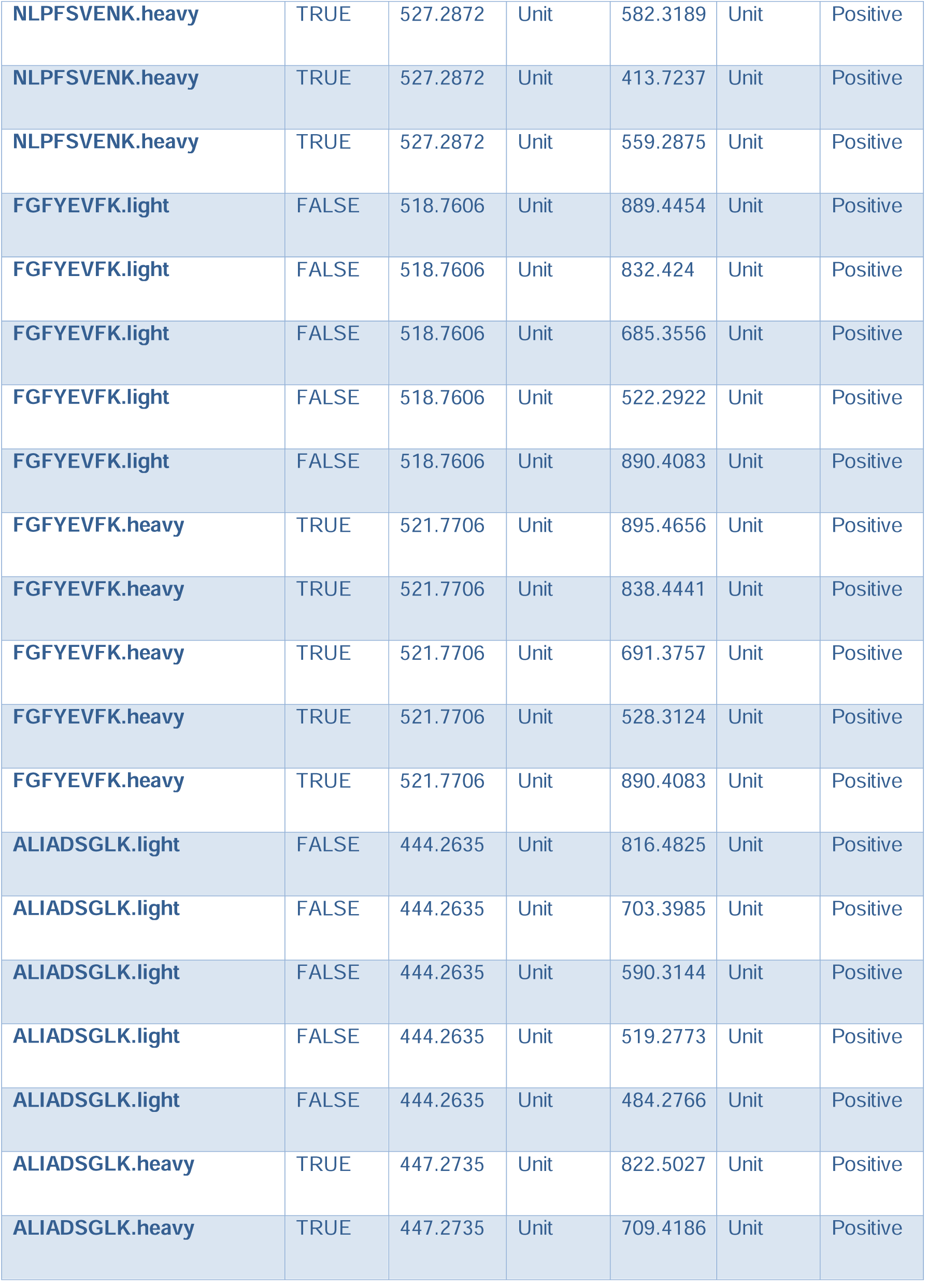
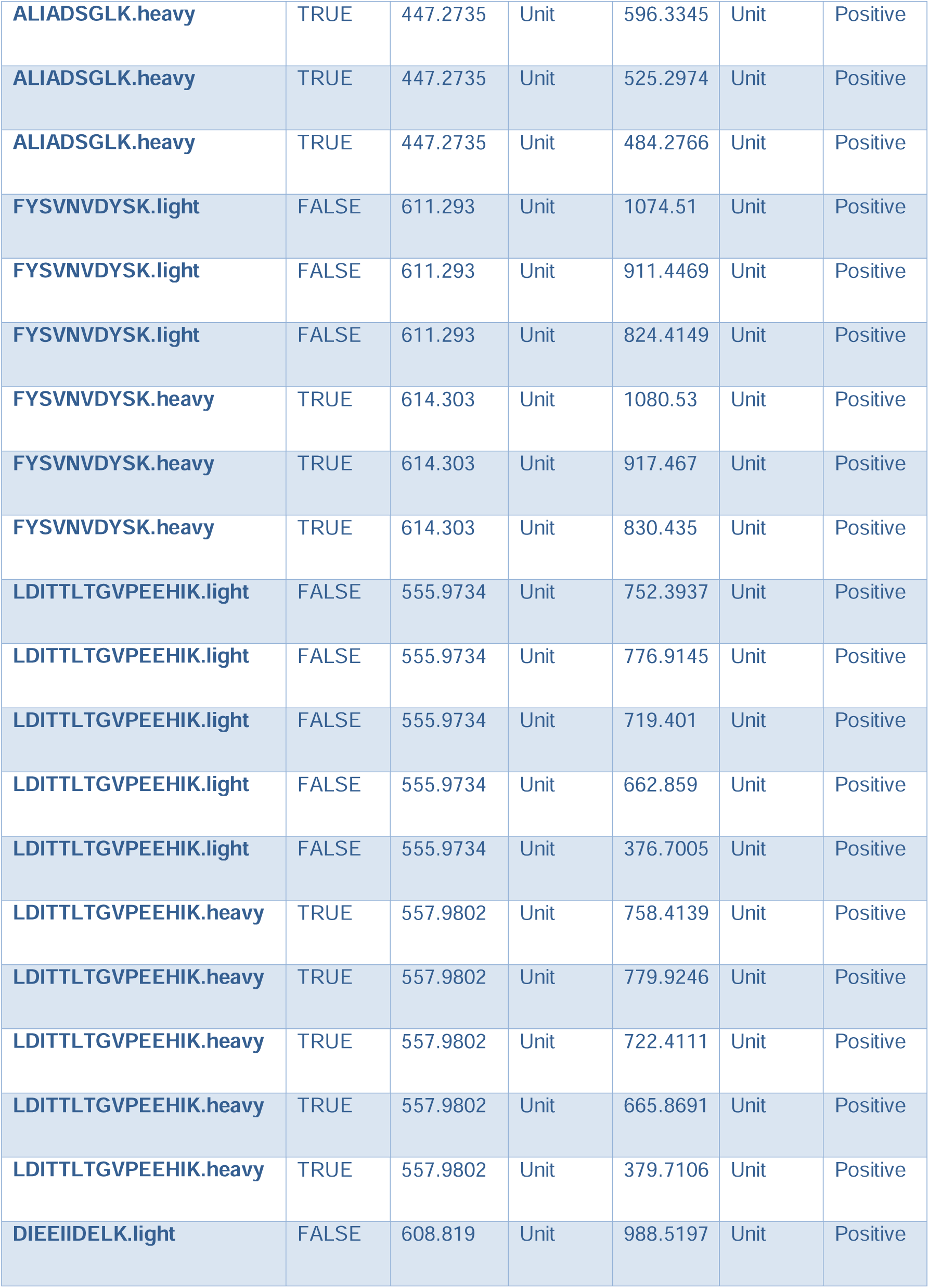
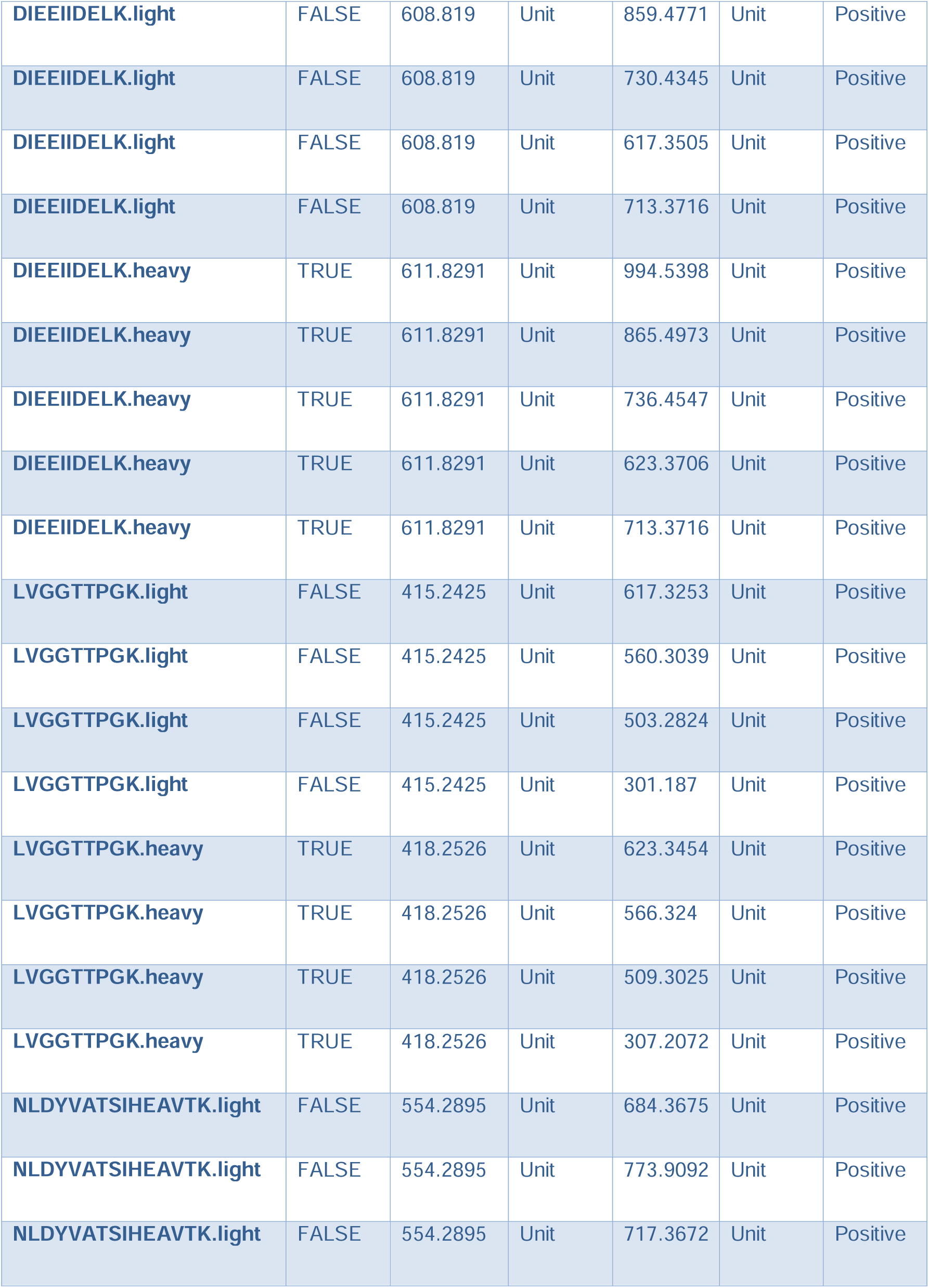
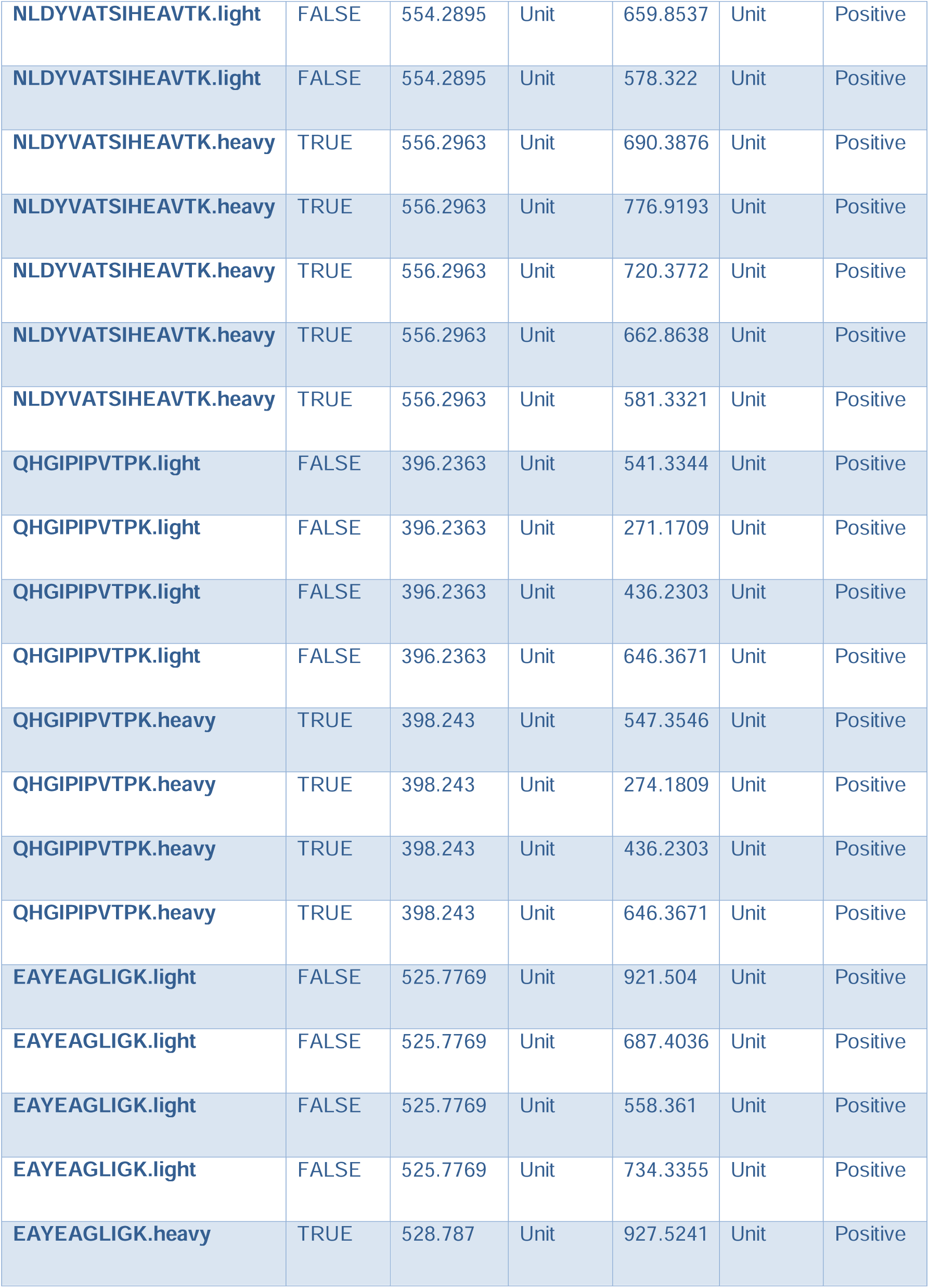
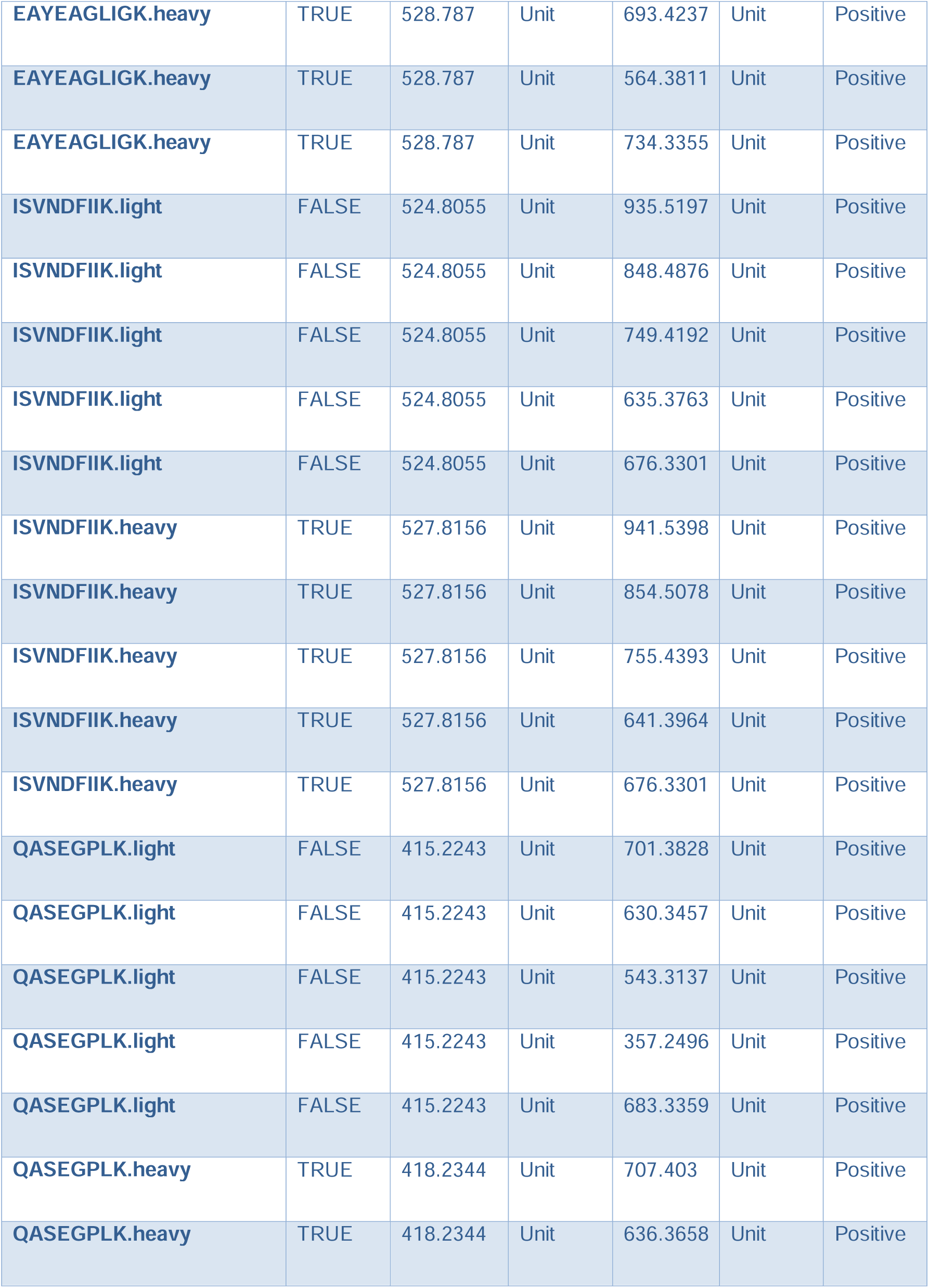
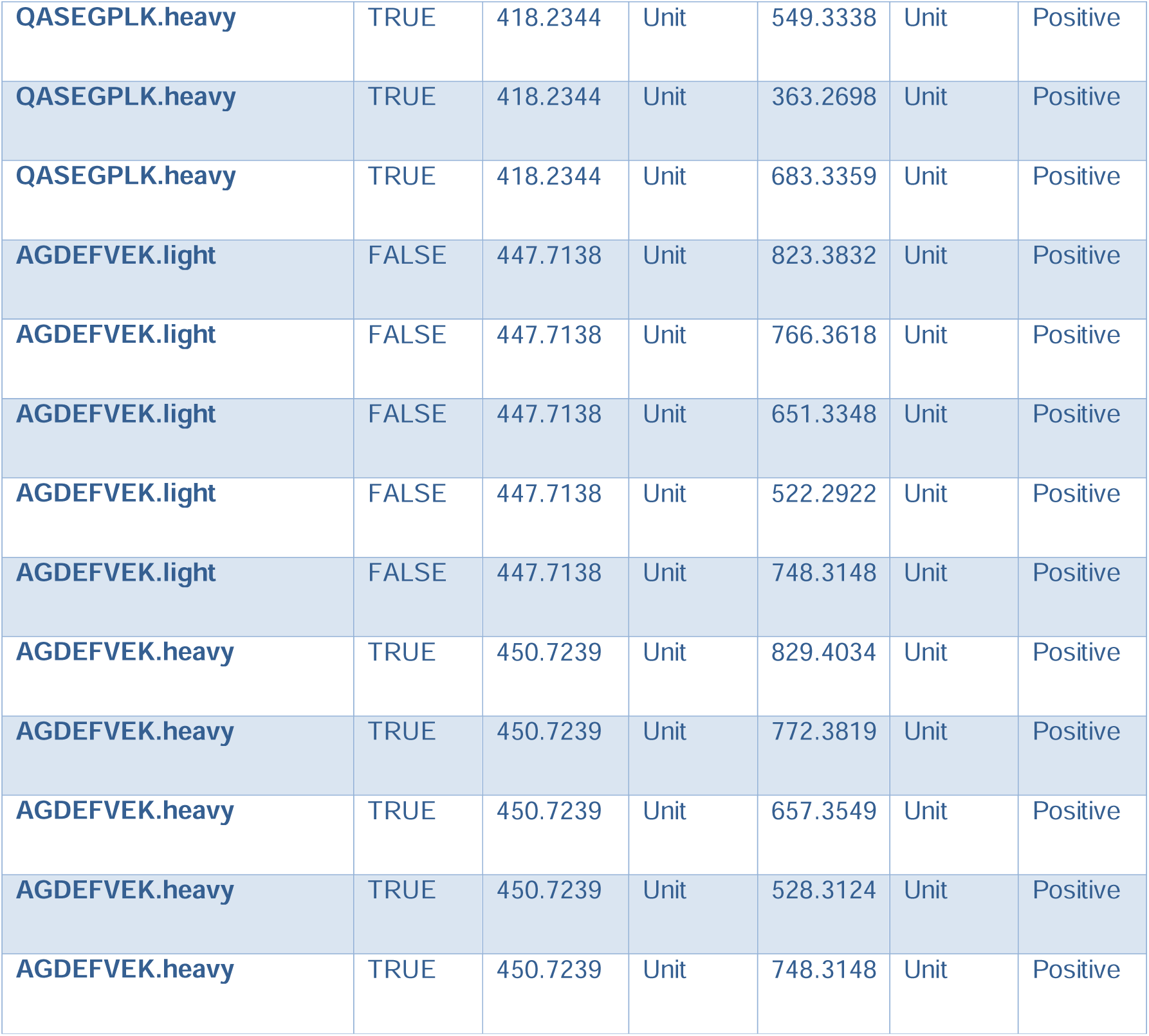
MRM transitions used to quantify the 14 selected proteins associated with various metabolic pathways.

