## Supplemental Data for "SysQuan: repurposing SILAC mice for the affordable absolute quantitation of the human proteome"

**Supplemental Data Table of Contents**

**Supplemental Data Figure S1.** Calibration curves to quantify 14 proteins associated with metabolic pathways

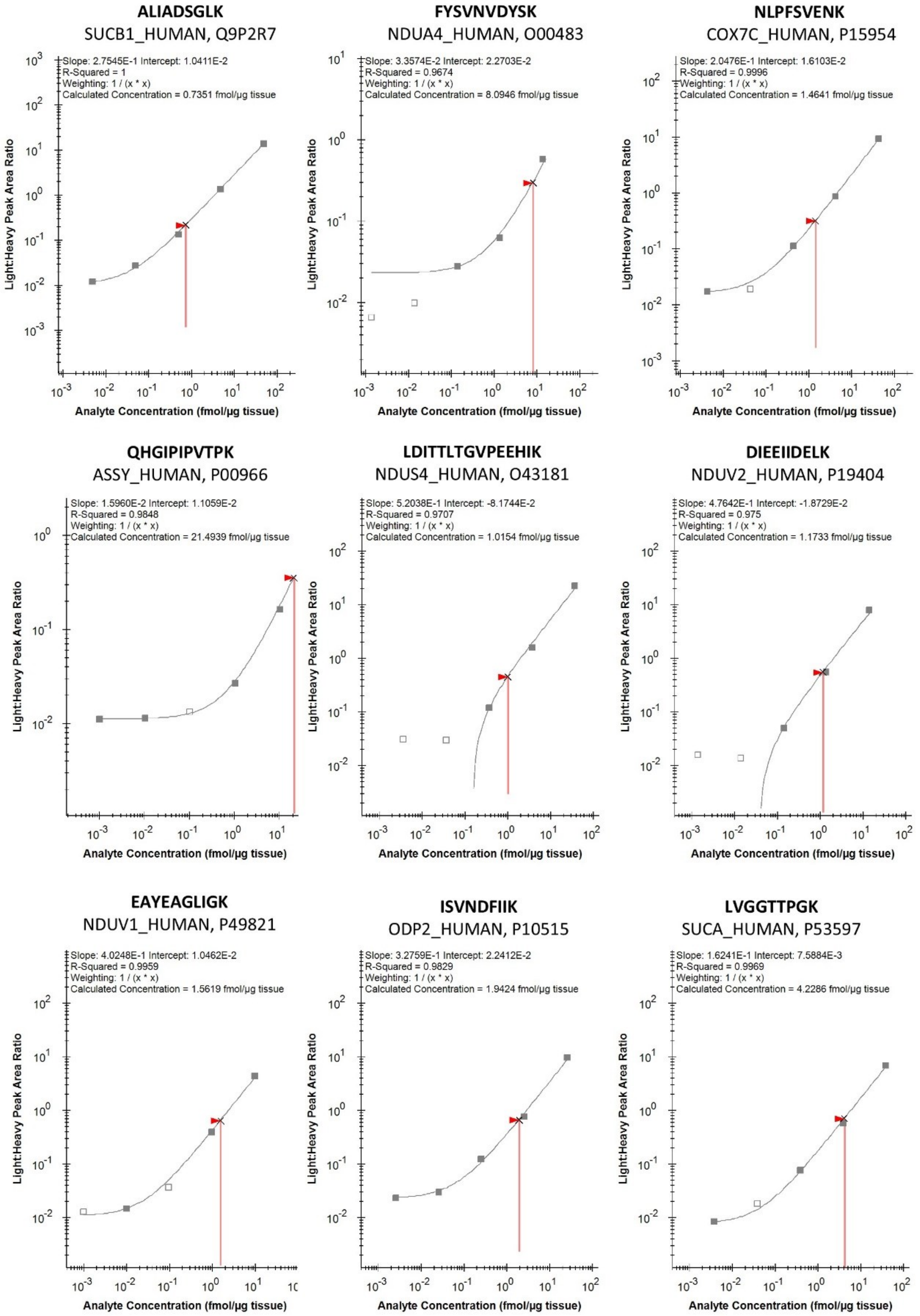

**QASEGPLK**
G3P_HUMAN, P04406

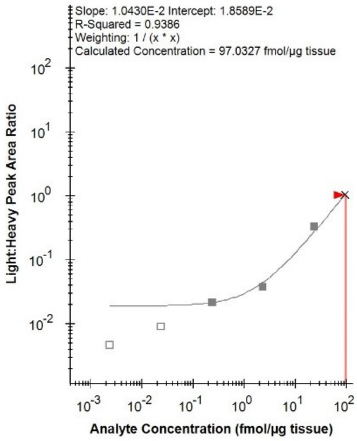

**FGFYEVFK**
MPCP_HUMAN, Q00325

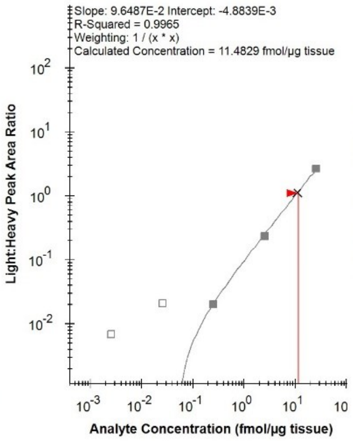

**YQQAGLPLIVLAGK**
ACOC_HUMAN, P21399

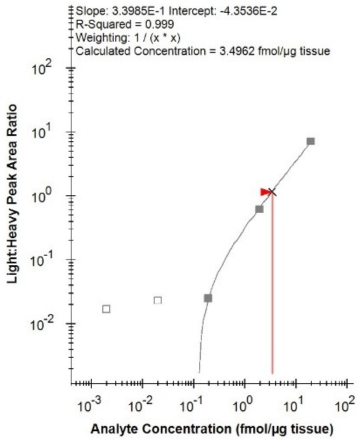

**AGDEFVEK**
HCDH_HUMAN, Q16836

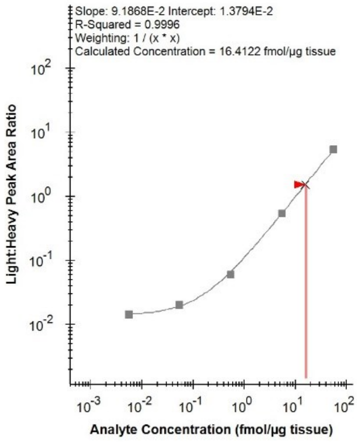

**NLDYVATSIHEAVTK**
AATC_HUMAN, P17174

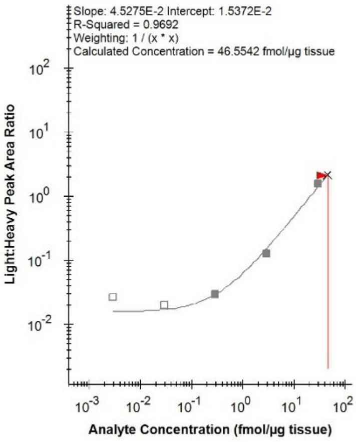

**Supplemental Data Table S1.** Proteins associated with various metabolic pathways for which we performed absolute quantification in human liver by double standards SILAC SIL and synthetic NAT

| Gene | ETC & Mitochondrial Function | TCA Cycle | Glycolysis | Amino Acid Metabolism | Beta-Oxidation | Mitochondrial Transport |
|---|---|---|---|---|---|---|
| ACO1 | | ✓ | | | | |
| ASS1 | | | | ✓ | | |
| COX7C | ✓ | | | | | |
| DLAT | | ✓ | | | | |
| GAPDH | | | ✓ | | | |
| GOT1 | | | | ✓ | | |
| HADH | | | | | ✓ | |
| NDUFA4 | ✓ | | | | | |
| NDUFS4 | ✓ | | | | | |
| NDUFV1 | ✓ | | | | | |
| NDUFV2 | ✓ | | | | | |
| SLC25A3 | | | | | | ✓ |
| SUCLA2 | | ✓ | | | | |
| SUCLG1 | | ✓ | | | | |

**Supplemental Data Table S2.** MRM transitions used to quantify the 14 selected proteins associated with various metabolic pathways

| Compound Name | ISTD? | Precursor Ion | MS1 Res | Product Ion | MS2 Res | Polarity |
|---|---|---|---|---|---|---|
| **YQQAGLPLIVLAGK.light** | FALSE | 490.9608 | Unit | 810.5448 | Unit | Positive |
| **YQQAGLPLIVLAGK.light** | FALSE | 490.9608 | Unit | 713.492 | Unit | Positive |
| **YQQAGLPLIVLAGK.light** | FALSE | 490.9608 | Unit | 600.4079 | Unit | Positive |
| **YQQAGLPLIVLAGK.light** | FALSE | 490.9608 | Unit | 405.776 | Unit | Positive |
| **YQQAGLPLIVLAGK.light** | FALSE | 490.9608 | Unit | 661.3304 | Unit | Positive |
| **YQQAGLPLIVLAGK.heavy** | TRUE | 492.9675 | Unit | 816.5649 | Unit | Positive |
| **YQQAGLPLIVLAGK.heavy** | TRUE | 492.9675 | Unit | 719.5121 | Unit | Positive |
| **YQQAGLPLIVLAGK.heavy** | TRUE | 492.9675 | Unit | 606.4281 | Unit | Positive |
| **YQQAGLPLIVLAGK.heavy** | TRUE | 492.9675 | Unit | 408.7861 | Unit | Positive |
| **YQQAGLPLIVLAGK.heavy** | TRUE | 492.9675 | Unit | 661.3304 | Unit | Positive |
| **NLPFSVENK.light** | FALSE | 524.2771 | Unit | 820.4199 | Unit | Positive |
| **NLPFSVENK.light** | FALSE | 524.2771 | Unit | 723.3672 | Unit | Positive |
| **NLPFSVENK.light** | FALSE | 524.2771 | Unit | 576.2988 | Unit | Positive |
| **NLPFSVENK.light** | FALSE | 524.2771 | Unit | 410.7136 | Unit | Positive |
| **NLPFSVENK.light** | FALSE | 524.2771 | Unit | 559.2875 | Unit | Positive |
| **NLPFSVENK.heavy** | TRUE | 527.2872 | Unit | 826.4401 | Unit | Positive |
| **NLPFSVENK.heavy** | TRUE | 527.2872 | Unit | 729.3873 | Unit | Positive |
| **NLPFSVENK.heavy** | TRUE | 527.2872 | Unit | 582.3189 | Unit | Positive |
| **NLPFSVENK.heavy** | TRUE | 527.2872 | Unit | 413.7237 | Unit | Positive |
| **NLPFSVENK.heavy** | TRUE | 527.2872 | Unit | 559.2875 | Unit | Positive |
| **FGFYEVFK.light** | FALSE | 518.7606 | Unit | 889.4454 | Unit | Positive |
| **FGFYEVFK.light** | FALSE | 518.7606 | Unit | 832.424 | Unit | Positive |
| **FGFYEVFK.light** | FALSE | 518.7606 | Unit | 685.3556 | Unit | Positive |
| **FGFYEVFK.light** | FALSE | 518.7606 | Unit | 522.2922 | Unit | Positive |
| **FGFYEVFK.light** | FALSE | 518.7606 | Unit | 890.4083 | Unit | Positive |
| **FGFYEVFK.heavy** | TRUE | 521.7706 | Unit | 895.4656 | Unit | Positive |
| **FGFYEVFK.heavy** | TRUE | 521.7706 | Unit | 838.4441 | Unit | Positive |
| **FGFYEVFK.heavy** | TRUE | 521.7706 | Unit | 691.3757 | Unit | Positive |
| **FGFYEVFK.heavy** | TRUE | 521.7706 | Unit | 528.3124 | Unit | Positive |
| **FGFYEVFK.heavy** | TRUE | 521.7706 | Unit | 890.4083 | Unit | Positive |
| **ALIADSGLK.light** | FALSE | 444.2635 | Unit | 816.4825 | Unit | Positive |
| **ALIADSGLK.light** | FALSE | 444.2635 | Unit | 703.3985 | Unit | Positive |
| **ALIADSGLK.light** | FALSE | 444.2635 | Unit | 590.3144 | Unit | Positive |
| **ALIADSGLK.light** | FALSE | 444.2635 | Unit | 519.2773 | Unit | Positive |
| **ALIADSGLK.light** | FALSE | 444.2635 | Unit | 484.2766 | Unit | Positive |
| **ALIADSGLK.heavy** | TRUE | 447.2735 | Unit | 822.5027 | Unit | Positive |
| **ALIADSGLK.heavy** | TRUE | 447.2735 | Unit | 709.4186 | Unit | Positive |
| **ALIADSGLK.heavy** | TRUE | 447.2735 | Unit | 596.3345 | Unit | Positive |
| **ALIADSGLK.heavy** | TRUE | 447.2735 | Unit | 525.2974 | Unit | Positive |
| **ALIADSGLK.heavy** | TRUE | 447.2735 | Unit | 484.2766 | Unit | Positive |
| **FYSVNVDYSK.light** | FALSE | 611.293 | Unit | 1074.51 | Unit | Positive |

| | | | | | | |
|---|---|---|---|---|---|---|
| **FYSVNVDYSK.light** | FALSE | 611.293 | Unit | 911.4469 | Unit | Positive |
| **FYSVNVDYSK.light** | FALSE | 611.293 | Unit | 824.4149 | Unit | Positive |
| **FYSVNVDYSK.heavy** | TRUE | 614.303 | Unit | 1080.53 | Unit | Positive |
| **FYSVNVDYSK.heavy** | TRUE | 614.303 | Unit | 917.467 | Unit | Positive |
| **FYSVNVDYSK.heavy** | TRUE | 614.303 | Unit | 830.435 | Unit | Positive |
| **LDITTLTGVPEEHIK.light** | FALSE | 555.9734 | Unit | 752.3937 | Unit | Positive |
| **LDITTLTGVPEEHIK.light** | FALSE | 555.9734 | Unit | 776.9145 | Unit | Positive |
| **LDITTLTGVPEEHIK.light** | FALSE | 555.9734 | Unit | 719.401 | Unit | Positive |
| **LDITTLTGVPEEHIK.light** | FALSE | 555.9734 | Unit | 662.859 | Unit | Positive |
| **LDITTLTGVPEEHIK.light** | FALSE | 555.9734 | Unit | 376.7005 | Unit | Positive |
| **LDITTLTGVPEEHIK.heavy** | TRUE | 557.9802 | Unit | 758.4139 | Unit | Positive |
| **LDITTLTGVPEEHIK.heavy** | TRUE | 557.9802 | Unit | 779.9246 | Unit | Positive |
| **LDITTLTGVPEEHIK.heavy** | TRUE | 557.9802 | Unit | 722.4111 | Unit | Positive |
| **LDITTLTGVPEEHIK.heavy** | TRUE | 557.9802 | Unit | 665.8691 | Unit | Positive |
| **LDITTLTGVPEEHIK.heavy** | TRUE | 557.9802 | Unit | 379.7106 | Unit | Positive |
| **DIEEIIDELK.light** | FALSE | 608.819 | Unit | 988.5197 | Unit | Positive |
| **DIEEIIDELK.light** | FALSE | 608.819 | Unit | 859.4771 | Unit | Positive |
| **DIEEIIDELK.light** | FALSE | 608.819 | Unit | 730.4345 | Unit | Positive |
| **DIEEIIDELK.light** | FALSE | 608.819 | Unit | 617.3505 | Unit | Positive |
| **DIEEIIDELK.light** | FALSE | 608.819 | Unit | 713.3716 | Unit | Positive |
| **DIEEIIDELK.heavy** | TRUE | 611.8291 | Unit | 994.5398 | Unit | Positive |
| **DIEEIIDELK.heavy** | TRUE | 611.8291 | Unit | 865.4973 | Unit | Positive |
| **DIEEIIDELK.heavy** | TRUE | 611.8291 | Unit | 736.4547 | Unit | Positive |
| **DIEEIIDELK.heavy** | TRUE | 611.8291 | Unit | 623.3706 | Unit | Positive |
| **DIEEIIDELK.heavy** | TRUE | 611.8291 | Unit | 713.3716 | Unit | Positive |
| **LVGGTTPGK.light** | FALSE | 415.2425 | Unit | 617.3253 | Unit | Positive |
| **LVGGTTPGK.light** | FALSE | 415.2425 | Unit | 560.3039 | Unit | Positive |
| **LVGGTTPGK.light** | FALSE | 415.2425 | Unit | 503.2824 | Unit | Positive |
| **LVGGTTPGK.light** | FALSE | 415.2425 | Unit | 301.187 | Unit | Positive |
| **LVGGTTPGK.heavy** | TRUE | 418.2526 | Unit | 623.3454 | Unit | Positive |
| **LVGGTTPGK.heavy** | TRUE | 418.2526 | Unit | 566.324 | Unit | Positive |
| **LVGGTTPGK.heavy** | TRUE | 418.2526 | Unit | 509.3025 | Unit | Positive |
| **LVGGTTPGK.heavy** | TRUE | 418.2526 | Unit | 307.2072 | Unit | Positive |
| **NLDYVATSIHEAVTK.light** | FALSE | 554.2895 | Unit | 684.3675 | Unit | Positive |
| **NLDYVATSIHEAVTK.light** | FALSE | 554.2895 | Unit | 773.9092 | Unit | Positive |
| **NLDYVATSIHEAVTK.light** | FALSE | 554.2895 | Unit | 717.3672 | Unit | Positive |
| **NLDYVATSIHEAVTK.light** | FALSE | 554.2895 | Unit | 659.8537 | Unit | Positive |
| **NLDYVATSIHEAVTK.light** | FALSE | 554.2895 | Unit | 578.322 | Unit | Positive |
| **NLDYVATSIHEAVTK.heavy** | TRUE | 556.2963 | Unit | 690.3876 | Unit | Positive |
| **NLDYVATSIHEAVTK.heavy** | TRUE | 556.2963 | Unit | 776.9193 | Unit | Positive |
| **NLDYVATSIHEAVTK.heavy** | TRUE | 556.2963 | Unit | 720.3772 | Unit | Positive |
| **NLDYVATSIHEAVTK.heavy** | TRUE | 556.2963 | Unit | 662.8638 | Unit | Positive |
| **NLDYVATSIHEAVTK.heavy** | TRUE | 556.2963 | Unit | 581.3321 | Unit | Positive |
| **QHGIPIPVTPK.light** | FALSE | 396.2363 | Unit | 541.3344 | Unit | Positive |
| **QHGIPIPVTPK.light** | FALSE | 396.2363 | Unit | 271.1709 | Unit | Positive |
| **QHGIPIPVTPK.light** | FALSE | 396.2363 | Unit | 436.2303 | Unit | Positive |
| **QHGIPIPVTPK.light** | FALSE | 396.2363 | Unit | 646.3671 | Unit | Positive |

| | | | | | | |
|---|---|---|---|---|---|---|
| **QHGIPIPVTPK.heavy** | TRUE | 398.243 | Unit | 547.3546 | Unit | Positive |
| **QHGIPIPVTPK.heavy** | TRUE | 398.243 | Unit | 274.1809 | Unit | Positive |
| **QHGIPIPVTPK.heavy** | TRUE | 398.243 | Unit | 436.2303 | Unit | Positive |
| **QHGIPIPVTPK.heavy** | TRUE | 398.243 | Unit | 646.3671 | Unit | Positive |
| **EAYEAGLIGK.light** | FALSE | 525.7769 | Unit | 921.504 | Unit | Positive |
| **EAYEAGLIGK.light** | FALSE | 525.7769 | Unit | 687.4036 | Unit | Positive |
| **EAYEAGLIGK.light** | FALSE | 525.7769 | Unit | 558.361 | Unit | Positive |
| **EAYEAGLIGK.light** | FALSE | 525.7769 | Unit | 734.3355 | Unit | Positive |
| **EAYEAGLIGK.heavy** | TRUE | 528.787 | Unit | 927.5241 | Unit | Positive |
| **EAYEAGLIGK.heavy** | TRUE | 528.787 | Unit | 693.4237 | Unit | Positive |
| **EAYEAGLIGK.heavy** | TRUE | 528.787 | Unit | 564.3811 | Unit | Positive |
| **EAYEAGLIGK.heavy** | TRUE | 528.787 | Unit | 734.3355 | Unit | Positive |
| **ISVNDFIIK.light** | FALSE | 524.8055 | Unit | 935.5197 | Unit | Positive |
| **ISVNDFIIK.light** | FALSE | 524.8055 | Unit | 848.4876 | Unit | Positive |
| **ISVNDFIIK.light** | FALSE | 524.8055 | Unit | 749.4192 | Unit | Positive |
| **ISVNDFIIK.light** | FALSE | 524.8055 | Unit | 635.3763 | Unit | Positive |
| **ISVNDFIIK.light** | FALSE | 524.8055 | Unit | 676.3301 | Unit | Positive |
| **ISVNDFIIK.heavy** | TRUE | 527.8156 | Unit | 941.5398 | Unit | Positive |
| **ISVNDFIIK.heavy** | TRUE | 527.8156 | Unit | 854.5078 | Unit | Positive |
| **ISVNDFIIK.heavy** | TRUE | 527.8156 | Unit | 755.4393 | Unit | Positive |
| **ISVNDFIIK.heavy** | TRUE | 527.8156 | Unit | 641.3964 | Unit | Positive |
| **ISVNDFIIK.heavy** | TRUE | 527.8156 | Unit | 676.3301 | Unit | Positive |
| **QASEGPLK.light** | FALSE | 415.2243 | Unit | 701.3828 | Unit | Positive |
| **QASEGPLK.light** | FALSE | 415.2243 | Unit | 630.3457 | Unit | Positive |
| **QASEGPLK.light** | FALSE | 415.2243 | Unit | 543.3137 | Unit | Positive |
| **QASEGPLK.light** | FALSE | 415.2243 | Unit | 357.2496 | Unit | Positive |
| **QASEGPLK.light** | FALSE | 415.2243 | Unit | 683.3359 | Unit | Positive |
| **QASEGPLK.heavy** | TRUE | 418.2344 | Unit | 707.403 | Unit | Positive |
| **QASEGPLK.heavy** | TRUE | 418.2344 | Unit | 636.3658 | Unit | Positive |
| **QASEGPLK.heavy** | TRUE | 418.2344 | Unit | 549.3338 | Unit | Positive |
| **QASEGPLK.heavy** | TRUE | 418.2344 | Unit | 363.2698 | Unit | Positive |
| **QASEGPLK.heavy** | TRUE | 418.2344 | Unit | 683.3359 | Unit | Positive |
| **AGDEFVEK.light** | FALSE | 447.7138 | Unit | 823.3832 | Unit | Positive |
| **AGDEFVEK.light** | FALSE | 447.7138 | Unit | 766.3618 | Unit | Positive |
| **AGDEFVEK.light** | FALSE | 447.7138 | Unit | 651.3348 | Unit | Positive |
| **AGDEFVEK.light** | FALSE | 447.7138 | Unit | 522.2922 | Unit | Positive |
| **AGDEFVEK.light** | FALSE | 447.7138 | Unit | 748.3148 | Unit | Positive |
| **AGDEFVEK.heavy** | TRUE | 450.7239 | Unit | 829.4034 | Unit | Positive |
| **AGDEFVEK.heavy** | TRUE | 450.7239 | Unit | 772.3819 | Unit | Positive |
| **AGDEFVEK.heavy** | TRUE | 450.7239 | Unit | 657.3549 | Unit | Positive |
| **AGDEFVEK.heavy** | TRUE | 450.7239 | Unit | 528.3124 | Unit | Positive |
| **AGDEFVEK.heavy** | TRUE | 450.7239 | Unit | 748.3148 | Unit | Positive |
